# Contrasting ecological communities in rice paddy fields under conventional and no-fertilizer farming practices

**DOI:** 10.1101/2025.08.12.669982

**Authors:** Masayuki Ushio, Hiroki Saito, Yoshiko Shimono, Kazunori Okada, Rika Ozawa, Kaori Shiojiri, Junji Takabayashi

**Affiliations:** Department of Ocean Science, The Hong Kong University of Science and Technology, Clear Water Bay, Kowloon, Hong Kong SAR; Tropical Agriculture Research Front, Japan International Research Center for Agricultural Sciences, 1091-1 Maezato-Kawarabaru, Ishigaki, Okinawa 907-0002, Japan; Graduate School of Agriculture, Kyoto University, Oiwake-cho, Kitashirakawa, Sakyo-ku, Kyoto 606-8502, Japan; Agro-Biotechnology Research Center, the University of Tokyo, 113-8657, Tokyo, Japan; Center for Ecological Research, Kyoto University, 2-509-3 Hirano, Otsu, Shiga 520-2113, Japan; Faculty of Agriculture, Ryukoku University, 1-5 Yokotani, Seta Oe-cho, Otsu, Shiga 520-2194, Japan

**Keywords:** Agricultural system, Ecological monitoring, Ecological variables, Environmental DNA, Rice, Time-series analysis, Unified information-theoretic causality

## Abstract

Global crop production is essential for providing energy and nutrients to humans, but agricultural systems contribute substantially to environmental issues, making it crucial to find sustainable methods for crop production. In this study, we conducted extensive field monitoring of two rice paddy fields in Kyoto Prefecture, Japan, where a rice variety, *Hinohikari*, was cultivated to investigate how ecological communities influence rice growth and yields under two contrasting farming practices: conventional farming and no-fertilizer farming. We had three objectives: (1) to monitor rice growth and ecological dynamics to create a comprehensive ecological time series using quantitative environmental DNA metabarcoding and complementary monitoring methods, (2) to identify ecological variables that causally affect rice performance using nonlinear time series analysis, and (3) to examine the effects of the two farming practices on ecological variables and rice performance. As expected, the no-fertilizer paddy field showed lower rice growth and yields but still produced 40–50% of the yields of the conventional field. Ecological monitoring revealed contrasting ecological communities between the two fields, particularly among plant species on the paddy ridges and microbial taxa. Twenty-five taxa had statistically clear causal influences on rice performance. While the per-abundance influences of the causal taxa were largely similar in both paddy fields, their abundances were different, contributing to the differences in the overall effects of these taxa on the rice performance. These findings suggest that the abundance of causal taxa, along with nutrient conditions, may have driven differences in rice growth between the two paddy fields.

## 1. INTRODUCTION

Global production of crops provides energy and nutrients for humans (Godfray et al., 2010), and rice (*Oryza sativa*) is one of the most important crops, along with maize, wheat, sugar cane, and oil palm, comprising 8% of world crop production (FAO, 2024). As with many other crops, improving yields and production efficiency such as resource use efficiency and pathogen resistance is one of the major concerns in rice production (e.g., Zhang, 2007), which is essential for the sustainability of human society. However, agriculture systems are significant contributors to greenhouse gas emissions and other environmental issues such as nutrient runoff as nonpoint source pollution (Hua et al., 2017; Menegat et al., 2022; Wang et al., 2014). Greenhouse gas emissions (CH_4_ and N_2_O) from paddy fields can affect the global climate (Qian et al., 2023), while nutrient runoff impacts surrounding ecosystems (Ko et al., 2021), both of which have deteriorating effects on the sustainability of the agriculture systems. Thus, finding ways to achieve sustainable rice crop production while minimizing environmental impacts is a key challenge in agricultural science.

Rice cultivation occurs in paddy fields, and rice growth, yields, and gene expression patterns can be influenced by various biotic and abiotic factors. Previous studies have examined how meteorological and endogenous variables (e.g., plant age and genotype) affect gene expression patterns (transcriptome dynamics) of rice in fluctuating field environments (Kashima et al., 2021; Nagano et al., 2012). These studies suggest that transcriptome dynamics are primarily driven by plant age and genotypes, and meteorological variables such as air temperature and solar radiation. Under these circumstances, genomics-based breeding, alongside advanced technologies such as high-throughput field-based phenotyping, is one of the most promising strategies to accommodate the complex interactions between rice and abiotic variables for improving rice yields and production efficiency (Wing et al., 2018).

Paddy fields are a habitat for a large number of species (e.g., more than 5000 species have been recorded in paddy fields in Japan; Natuhara, 2013), and ecological communities such as insect pests, aquatic animals, and microbial mutualists/pathogens significantly influence the rice performance (Ji et al., 2023; Li et al., 2025; Savary et al., 2019). For example, recurring outbreaks of an insect pest, brown planthopper, can cause serious physical and physiological damages to rice (Bottrell & Schoenly, 2012). Co-culturing rice with aquatic animals such as fish, crabs, crayfish, and turtles can improve the rice yields, reduce insect pests, and increase soil carbon and nitrogen contents (Ji et al., 2023; Yu et al., 2023). Also, fish can modify habitat structures in paddy fields through their foraging and swimming behaviour and excrement, and Li et al. (2025) showed that this effect could change the abundance of other key species such as earthworms and even paddy ecosystem functions, which could influence rice yields. Furthermore, not only organisms but also organic compounds such as green leaf volatiles (GLVs) can influence plant performance such as pathogen resistance (Matsui & Koeduka, 2016). The common GLVs (*E*)-2-hexenal and (*Z*)-3-hexenal induce defenses against pathogenic fungi (Matsui & Koeduka, 2016). Also, plant volatiles emitted by weeds on paddy ridges can influence rice physiological states and yields (Shiojiri et al., 2021). Thus, effects of ecological communities in a paddy field on rice should not be underestimated.

However, the dynamics of an ecological community are often challenging to monitor because of the large number of species (Natuhara, 2013) and the complex, state-dependent, and nonlinear nature of the dynamics (e.g., Hsieh et al., 2005). Furthermore, understanding the influence of ecological communities on rice performance, or interspecific interactions between rice and ecological communities, is even more difficult due to possible confounding factors under field conditions, despite the potential of such understanding for achieving sustainable agriculture (Ji et al., 2023; Toju et al., 2018). As a result, effects of only a relatively small number of species and/or variables on rice have been studied at one time in paddy fields (Ji et al., 2023; Li et al., 2025) and how a large number of ecological variables simultaneously affect the rice performance in paddy fields has remained unclear.

Environmental DNA (eDNA) analysis is one of the promising methods for overcoming the above-noted issues about comprehensive biodiversity monitoring under field conditions. eDNA refers to DNA directly extracted from environmental samples such as water and soil (Taberlet et al., 2018), which includes both microbial and macrobial DNA. Previous studies have shown that eDNA metabarcoding, an approach to comprehensively amplify and sequence marker regions of DNA belonging to a target taxa in an environmental sample, is a cost-and time-effective means to detect a large number of species (e.g., Deiner et al., 2016; Miya et al., 2015), and that eDNA-based community monitoring is especially informative when it is performed quantitatively (e.g., sequencing with internal spike-in DNAs: Ushio, 2022; Ushio, Murakami, et al., 2018). It enables tracking the detailed dynamics of a speciose ecological community (Ushio, 2022), and it can be even more effective when used with other traditional methods such as direct capture and visual census, as they can provide complementary information about biodiversity (He et al., 2023; Krol et al., 2019).

To investigate whether and how ecological variables influence rice performance under field conditions, we previously conducted eDNA-based, intensive monitoring of ecological community dynamics in five artificial rice plots (Ushio, 2022; Ushio et al., 2023). We collected water samples from artificial rice plots daily for 122 days and analyzed microbes and macrobes using a quantitative eDNA metabarcoding approach. To overcome another issue, that is, effects of confounding factors under fields condition (i.e., pseudo-correction or “mirage” correlation issue), we applied a time series-based causality detection method (Cenci et al., 2019; Deyle et al., 2016; Osada et al., 2023; Sugihara et al., 2012), to the time series, and identified species that affected the rice growth (i.e., an Oomycetes species, *Globisporangium nunn*, and a midge species, *Chironomus kiiensis*). Further, the effects of these two species on the rice growth and gene expression were validated using manipulative experiments. The findings of that study demonstrated that ecological community members indeed have effects on the rice growth performance and gene expression. However, the system used in Ushio et al. (2023) was artificial: small rice plots (about 1 m × 1m plot) and the farming practice was different from that used for real paddy fields; specifically, rice plants were grown in Wagner pots, irrigation water was maintained throughout the monitoring period, and no pesticides were applied. During the field monitoring, only eDNA metabarcoding was used to assess ecological communities, potentially overlooking important ecological variables such as flying insects, plants on paddy ridges, and specialized plant metabolites and their effects on rice in real paddy fields. In addition, time series-based causality detection methods have been actively developed (Runge et al., 2019; Suzuki et al., 2022) and several new methods have emerged, for example, the multiview distance regularized S-map (Chang et al., 2021), which enables more accurate quantification of interaction strengths. However, these new methods were not fully utilized in the previous studies (Ushio, 2022; Ushio et al., 2023).

In the present study, we conducted field monitoring of two rice paddy fields in Kyoto Prefecture, Japan, where rice variety *Hinohikari* was planted, to investigate whether and how ecological communities affect the rice growth and yields in real paddy fields. To study a wider range of environmental gradients, we targeted paddy fields with different farming practices and different backgrounds. The first one was a paddy field with conventional farming practice; namely, farmers applied basal nitrogen (N), phosphorus (P), and potassium (K) fertilizers and used pesticides to reduce plants on the paddy ridge. The second one was a paddy field with no fertilizer treatment. The farming practice of this field was unique, as it has not employed any basal fertilizers or pesticides since 1951 but still maintain rice yields equal to about 70–80% of the yields obtained with the conventional management practice (e.g., Okumura, 2002). How this “no-fertilizer” paddy field maintains its yield without nutrient inputs is unknown, but investigating the effects and dynamics of ecological communities may shed light on the mechanism of the maintenance of the yields. Given this background, the objectives of this study were: (1) to monitor rice growth and ecological dynamics to generate extensive time series using quantitative eDNA metabarcoding and other complementary monitoring approaches, (2) to identify ecological variables that causally influence rice performance and to quantify the causal effects using time series-based causality detection methods, and (3) to clarify the effects of the two contrasting management practices on the ecological variables and the rice performance, in real paddy fields.

## 2. METHODS

### 2.1 Study site

Our monitoring was performed at paddy fields in Uji, Kyoto, Japan (34°54′09″N, 135°46′29″E) from May to September in 2017 (Fig. 1a). Climate data during the field monitoring was retrieved from the Japan Meteorological Agency website (https://www.data.jma.go.jp/risk/obsdl/index.php, accessed on 1 November 2024; “daily mean values”, “Kyoto city”, and “2017” were selected in the webpage) (Fig. 1b–d), and the daily mean air temperatures were from 16.9°C to 31.2°C. The field monitoring was performed in two rice fields with different management practices: rice farming with fertilizer and pesticides (“conventional” farming; Fig. 1e) and that without fertilizer and pesticides (“no-fertilizer” farming; Fig. 1f). Hereafter, we refer to these fields as “conventional” and “no-fertilizer” paddy fields, respectively. The conventional paddy field occupies about 580 m^2^ and the no-fertilizer paddy field occupies about 945 m^2^ (Fig. S1a). They are separated by about 15 m (including a watercourse), and their irrigation waters were not mixed. While insect pests and a small fraction of microbes may migrate between the fields, the effects of the other treatment should be much smaller than those of the main treatment.

**Figure 1.**
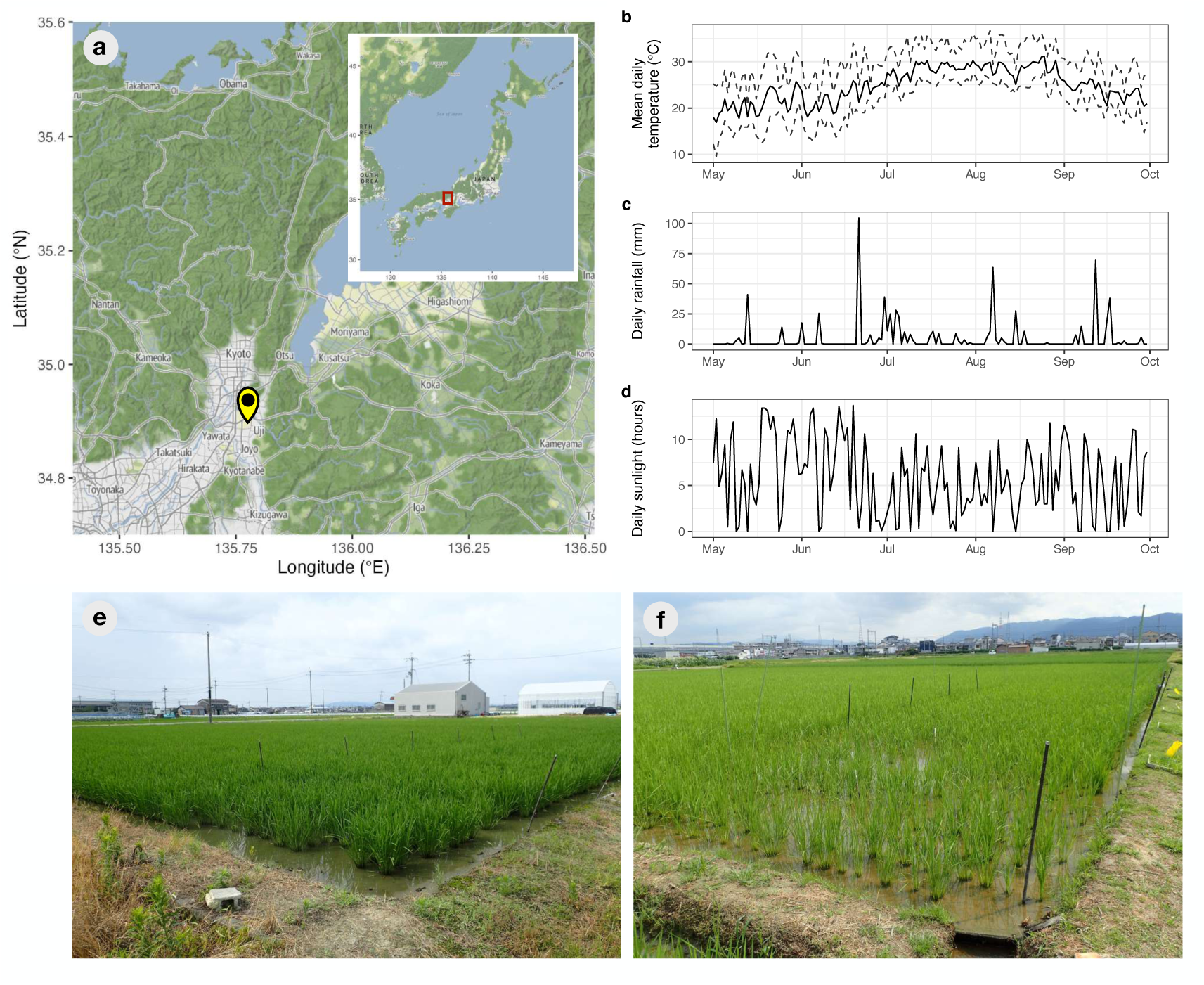
Study site, climate variables, and rice paddy fields with “conventional” and “no-fertilizer” farming practices. (**a**) Location of the study site. The yellow marker indicates the site, while the red rectangle in the inset highlights the study region (Maps created using the ggmap package of R; ©Stadia Maps, ©Stamen Design, ©OpenMapTiles and ©OpenStreetMap). (**b**) Mean daily air temperature (°C). (**c**) Daily rainfall (mm). (**d**) Daily sunlight hours. Climate data were obtained from the weather station in Kyoto City, managed by the Japan Meteorological Agency. (**e**) “Conventional” paddy field. (**f**) “No-fertilizer” paddy field. The yellow panel in the right side of the image was used for collecting insect samples. Note that the yellow panels were deployed in both fields. Field photographs were taken on 12 July 2017 (©M. Ushio).

In the conventional field, basal N-P-K fertilizer at 24–30 kg N ha^-1^ year^-1^ was added in May, and another 18 kg N ha^-1^ year^-1^ was subsequently added in mid-August. Three to four days after the rice planting, pesticide (ST Longet Floable, Sumitomo Chemical Company, Ltd., Tokyo, Japan) at 6,700 ml ha^-1^ was applied in the conventional field. The no-fertilizer paddy field has been maintained by a non-profit organization “Research Group on No Fertilizer and No Pesticide Farming” (https://muhiken.or.jp/). This paddy soil was initially located in Ritto, Shiga Prefecture (adjacent to Kyoto Prefecture; about 25 km from the location of the no-fertilizer paddy field), Japan, where farming without fertilizers or pesticides commenced in 1951 (https://muhiken.or.jp/ritto/, accessed as of 11 October 2025). In 2006, after farming in Ritto ceased, the no-fertilizer paddy soil was moved to Uji, Kyoto, where no-fertilizer rice farming has continued. Even without fertilizers, the no-fertilizer field maintained production of about 80% of the typical grain yield when rice variety *Beniasahi* was cultivated (Okumura, 2002). Furthermore, previous research has suggested differences in microbial community composition and nutrient uptake efficiency between the conventional and no-fertilizer fields (Kamata et al., 1991; Okumura, 2002). More detailed information about the no-fertilizer paddy field is provided in the Supplementary text.

### 2.2 Sampling design

In each paddy field, we conducted weekly monitoring of rice growth and ecological community dynamics from 19 May 2017 to 30 August 2017. The first monitoring was conducted on 19 May 2017 (Friday) before planting rice, var. *Hinohikari*, and the other monitoring was conducted in the morning (between 8:30 and 10:40) every Wednesday (see the sample metadata in Table S1). Although var. *Beniasahi* had previously been cultivated in the no-fertilizer paddy field, var. *Hinohikari* was selected for the present study to align with the variety grown by farmers in the conventional paddy field and to ensure consistency across the field types for evaluating ecological effects on rice growth. In total, we conducted the field survey 15 times, including the first one before planting the rice. The monitoring was ceased on 30 August because the rice field was drained in preparation for harvesting rice.

We set five rice monitoring points in each paddy field and five sampling points on each paddy ridge (Fig. S1a). In each paddy field, five 3 m × 3 m plots, which were 2 m away from the paddy ridge, were set to monitor the rice performance, including rice heights, SPAD values, and the number of rice stems (see the section “*2.3 Rice growth monitoring and yield survey*”). In each paddy ridge, we set sticky panels for collecting flying insects (Fig. S1b) and 50 cm × 50 cm quadrats (0.25 m^2^) for monitoring wild weeds (Fig. S1c) in each of five sampling points. At the same monitoring points in paddy fields, we collected rice leaves for analyzing plant specialized metabolites. Water samples for eDNA-based ecological community monitoring were collected from the rice monitoring points for quantitative monitoring of microbes and macrobes (see section “*2.4 Monitoring of ecological variables and communities*”). The rice was harvested on 29 September 2017, and the rice yield survey was conducted (Fig. S1d, e).

### 2.3 Rice growth monitoring and yield survey

Rice growth was monitored by measuring rice leaf height of target individuals at each monitoring point every week using a ruler (the highest leaf heights were measured). The average height of three individuals was used as a representative rice height of each monitoring point, and thus we had five representative values for each plot for each week. Leaf SPAD was also measured every week for the same leaf whose height was measured using a SPAD meter (SPAD-502Plus, KONICA-MINOLTA, Inc., Tokyo, Japan). The number of rice stems was also counted. After we harvested rice, we counted the number of fertile and sterile grains and measured their wet and dry weights as indices of rice yields.

### 2.4 Monitoring of ecological variables and communities

#### 2.4.1 Plant specialized metabolites

Rice leaves were collected for analyzing plant specialized metabolites and kept at –20°C during the transport to the laboratory. The volatile collection system using Gerstel-Twister (polydimethylsiloxane [PDMS] coated stir bar, film thickness 0.5 mm, 10 mm length, Gerstel GmbH & Co. KG) was used to collect volatile compounds emitted from the frozen rice leaves. Headspace volatiles emitted from four leaf pieces, each 4 cm long, were collected on two Twisters in a closed glass vial (30 ml) in a laboratory room (25 ± 1°C) for 3 hours.

Before starting the volatile collection, 0.1 µg of tridecane on a piece of filter paper was added to the glass bottle as an internal standard. The collected volatile compounds were analyzed by GC-MS with an HP-5MS capillary column as described previously (Rim et al., 2018). The peak area of each compound was normalized by dividing it by the peak area of the internal standard and the leaf weight (g). The normalized peak area was calculated as “relative peak area.”

Momilactone was quantified using rice leaf samples. Each leaf sample (10–400 mg FW) was immersed in 1 ml of 70% methanol and incubated at 4°C for 24 hours. A 2-μl aliquot of the extract was analyzed by LC–MS/MS (Sciex QTRAP 3200) with the following selected reaction monitoring (SRM) transitions (Riemann et al., 2013): momilactone A, m/z 315→271; momilactone B, m/z 331→269; phytocassanes A and E, m/z 317→299; phytocassane B, m/z 335→317; and phytocassane C, m/z 319→301. The total phytoalexin level was calculated as the sum of all compounds, reflecting the diterpenoid content. We detected momilactone A, momilactone B, phytocassane B, and phytocassane B3, and their concentrations were aggregated as “Diterpenoids (μg/g)” for statistical analysis.

#### 2.4.2 Insect collections

Yellow sticky traps (Horiver; 257 mm × 100 mm; Arysta LifeScience Corporation, Tokyo, Japan) were installed at the five sampling locations on the paddy ridges during the monitoring period. One Horiver trap was mounted on a plastic stick at each sampling point. The traps were replaced with new ones every week. The collected traps were examined under a stereomicroscope, and the numbers of thrips (Thysanoptera), leafhoppers (Cicadellidae), and other planthoppers (mainly Homoptera) were recorded.

#### 2.4.3 Plant cover survey on the ridge

Each 50 cm × 50 cm quadrat was subdivided into 25 cells of 10 cm × 10 cm by strings stretched across the frame (Fig. S1c). Photographs of the entire quadrat were taken from directly above the quadrat every week from June to August and the percentage cover of each major species was calculated as visual estimates from the photographs. On June 28 and July 26, direct vegetation surveys were also conducted on-site and confirmed that there were no discrepancies with the judgments made based on the photographs taken.

#### 2.4.4 Water sampling for eDNA analysis

Water samples, rather than soil samples, in each paddy field were collected for eDNA-based ecological community analysis to avoid soil sampling that could damage or stress the rice. The water was filtered using Sterivex filter cartridges (Merck Millipore, Darmstadt, Germany). Due to the high density of sediment in the rice paddy fields, the volume of water we could filter was relatively small (30–100 ml for each sampling point). Two types of filter cartridges were used to filter water samples: to detect microorganisms, φ0.22-µm Sterivex (SVGV010RS) filter cartridges that included zirconia beads inside were used to filter 30 ml of water (Ushio, 2019); and to detect macroorganisms, φ0.45-µm Sterivex (SVHV010RS) filter cartridges were used to filter 30–100 ml of water. After filtration, 2 ml of RNAlater solution (ThermoFisher Scientific, Waltham, Massachusetts, USA) were added to each filter cartridge to prevent DNA degradation during storage. The same volume of MilliQ water was filtered using the two types of filter cartridges as field negative controls. The filter cartridges were stored in a cooler box until they were taken back to the laboratory. They were stored at –20°C until further processing.

Although we conducted weekly water sampling from 19 May 2017 to 30 August 2017, water sampling was not possible for some sampling events because mid-season drainage was practiced in the rice fields to improve rice performance in August. In total, we collected 266 filter cartridges (118 field water samples × 2 filter cartridges + 15 field-level negative controls × 2 filter cartridges) from the two rice fields. Details about the water samples are available in Table S1.

### 2.5 Quantitative environmental DNA metabarcoding

#### 2.5.1 DNA extraction and library preparation for quantitative eDNA metabarcoding

Detailed protocols of the quantitative eDNA metabarcoding are described in the supplementary methods and Ushio (2022). Briefly, DNA was extracted and purified using a DNeasy Blood & Tissue kit (Qiagen, Hilden, Germany) (Ushio, 2019). After the purification, DNA was eluted using 100 µl of the elution buffer and stored at −20°C until further processing.

A two-step PCR approach was adopted for the library preparation. The first-round PCR (first PCR) was carried out with the internal standard DNAs to amplify metabarcoding regions using primers specific to prokaryotes (515F and 806R), eukaryotes (Euk_1391f and EukBr), fungi (ITS1-F-KYO1 and ITS2-KYO2) and animals (mlCOIintF and HCO2198). Sequences for primers and internal standard DNAs and primer references are available in the Supplementary methods. To monitor cross-contamination during the library preparation, 16 PCR-level negative controls with standard DNAs and 4 PCR-level negative controls without standard DNAs were included. In total, 153 DNA samples were analyzed for each type of filter cartridge (118 field water samples + 15 field-level negative controls + 20 PCR-level negative controls). The second-round PCR (second PCR) was carried out to append indices for different samples for sequencing. The DNA library was sequenced on the MiSeq platform (Illumina, San Diego, CA, USA) using the 250PE reagent kit for 16S, COI, and ITS, and the 150PE reagent kit for 18S.

#### 2.5.2 Sequence data processing

The raw sequence data were converted into FASTQ files using the bcl2fastq program provided by Illumina (bcl2fastq v2.18). The FASTQ files were then demultiplexed using the command implemented in Claident (http://www.claident.org) (Tanabe & Toju, 2013). Demultiplexed FASTQ files were then analyzed using the Amplicon Sequence Variant (ASV) method implemented in the “dada2” package (Callahan et al., 2016) of R (R Core Team, 2024). ASVs detected in the three experiments were merged and clustered into OTUs at 97% similarity using the DECIPHER package of R (Wright, 2016), which converted the ASV-sample matrix into the OTU-sample matrix. We converted ASVs to OTUs because ASVs often detected intraspecific variations and the focus of this experiment was community compositions at the species level (not at the intra-species level). Taxonomic identification was performed using Claident v0.9.2024.03.04 (Tanabe & Toju, 2013).

For all analyses after this subsection, the free statistical environment R was used (R Core Team, 2024). The procedure used to estimate DNA copy numbers consisted of two parts, following previous studies (Ushio, 2019; Ushio, Murakami, et al., 2018). Briefly, we performed (i) linear regression analysis to examine the relationship between sequence reads and the copy numbers of the internal standard DNAs for each sample, and (ii) conversion of sequence reads of non-standard DNAs to estimate the copy numbers using the result of the linear regression for each sample. The regression equation was: Sequence reads = sample-specific regression slope × the number of standard DNA copies [/µl]. Then, the estimated copy numbers per µl extracted DNA (copies/µl) were converted to DNA copy numbers per ml water in the rice plot (copies/ml water). After the conversion to DNA copy numbers, rare OTUs (their concentrations were lower than half of the minimum concentration of standard DNA: 1,250 copies/ml for 16S, 12.5 copies/ml for 18S, 1.0625 copies/ml for COI, and 10 copies/ml for ITS) and infrequently detected OTUs (detected in less than 10% of total monitoring events) were removed to mitigate the effects of sequencing errors.

### 2.6 Statistical analysis and visualization of ecological community dynamics

The monitoring data on rice performance and ecological communities were initially visualized through time-series plots created with the ggplot2 package in R (Wickham, 2009). Subsequently, the effects of time (sampling date) and farming practices were analyzed using generalized additive mixed models (GAMM) for rice performance, plant specialized metabolites, insect abundance, plant cover, and eDNA abundance and diversity, incorporating random effects for rice individuals (when applicable) and sampling locations. GAMM analysis was performed using the “mgcv” package in R (Wood, 2004). The influence of farming practices on the ecological variables affecting rice performance (see the next section) was assessed using linear mixed models (LMM) with random effects for sampling locations, utilizing the “nlme” package in R (Pinheiro et al., 2009). Additionally, the effect of management practices on rice yields was evaluated using analysis of variance (ANOVA).

Non-metric Multi-dimensional Scaling (NMDS) was performed to visualize the differences in plant cover on the ridge, plant specialized metabolites, and ecological communities between the two paddy fields. The `adonis2()` function was utilized to assess whether time (i.e., the number of weeks) and farming practices had statistically clear effects. In the eDNA community composition data, taxa that differentiated the two paddy fields were identified using the `envfit()` function. The “vegan” package in R (Oksanen et al., 2008) was employed for both NMDS and `adonis2()` analyses. Throughout the statistical analysis, we use the phrase “statistically clear” instead of “significant” in this study to clarify the meaning of the word “significant” by following Dushoff et al. (2019).

### 2.7 Time series analysis-based causality detection and quantification of influences from ecological variables on rice growth

To identify factors influencing the rice performance, we quantified information flow between the rice performance and ecological variables by the unified information-theoretic causality (UIC) method (Osada et al., 2023) implemented in the “rUIC” v0.9.13 package (Osada & Ushio, 2021) of R. UIC tests the statistical clarity of information flow between variables in terms of TE (Schreiber, 2000) computed by nearest neighbor regression based on time-delay embedding of explanatory variables (i.e., cross mapping; Sugihara et al. 2012). In contrast to the standard method used to measure TE, UIC quantifies information flow as follows:

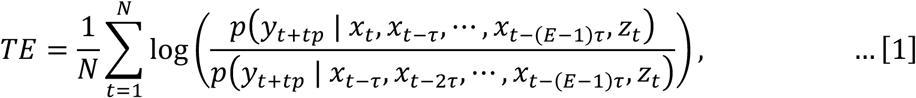

where *x*, *y*, and *z* represent an effect variable, a potential causal variable, and a conditional variable (if available), respectively. 𝑝(𝐴|𝐵) represents conditional probability: the probability of *A* conditioned on *B*. *t*, *tp*, *τ*, and *E* represent the time index, time step (time-lag), a unit of time-lag, and the optimal embedding dimension, respectively. *N* is the total number of points in the reconstructed state space (this is equivalent to the total number of time points – the optimal embedding dimension + 1). For example, if *tp* = –1 in Eqn. [1], UIC tests the causal effect from *yt*–1 to *xt*. Optimal *E* was selected using a TE version of simplex projection (Sugihara & May, 1990) implemented in the rUIC package. In the present study, the causality time-lag (*tp*) up to –2 (equivalent to two-week time-lag) was tested. Statistical clarity was tested by bootstrapping data after embedding (the statistical clarity levels for each time-lag were set to 0.05).

Because most of our time-series data exhibited a clear trend (e.g., rice height showed an increasing trend until August), we took the first-difference of the original time-series to make the time series stationary for the UIC analysis. To determine the variables that causally influence rice performance, we quantified TE between the indices of rice performance (i.e., growth rate, SPAD, and the number of stems) and climate and ecological variables. Given the large number of statistical tests required (the total number of UIC tests = [rice performance indices] × tp [0, –1, or –2] × [climate and ecological variables]), we calculated the false discovery rate (FDR) (Benjamini & Hochberg, 1995) for each combination of the rice performance indices and causal (climate and ecological) variables. We judged that a variable has a statistically clear causal effect on rice performance if the FDR is below 0.05. We first performed the UIC analysis between the climate variables and the rice performance. Then, we included climate variables that have statistically clear effects on the rice performance in the embedding in the UIC analysis to reduce the possibility of misidentification of causality from ecological variables on the rice performance due to the strong effect of climate variables.

### 2.8 Nonlinear time series analysis to quantify interaction strengths

We quantified the effects of ecological variables on the rice performance using an improved version of the sequential locally weighted global linear map (S-map) (Sugihara, 1994), called the multiview distance regularized S-map (MDR S-map) (Chang et al., 2021). Consider a system that has *E* different interacting variables and assume that the state space at time *t* is given by 𝑿_𝑡_ = {𝑥_1,𝑡_, 𝑥_2,𝑡_, … , 𝑥_𝐸,𝑡_}. For each target time point 𝑡^∗^, the S-map method produces a local linear model that predicts the future value 𝑥_1,_*_𝑡_*_∗+𝑝_ from the multivariate reconstructed state space vector 𝑿_𝑡_∗ = {𝑥_1,𝑡_∗, 𝑥_2,𝑡_∗, … , 𝑥_𝐸,𝑡_∗}. That is,

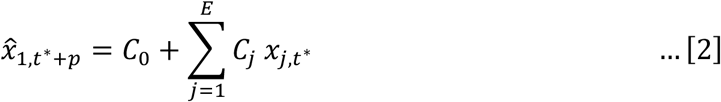

where ̂*𝑥*_1,_*_𝑡_*_∗+𝑝_ is a predicted value of 𝑥_1_ at time 𝑡^∗^ + 𝑝, 𝐶_j_ is a regression coefficient and interpreted as interaction strength (or called S-map coefficient), and 𝐶_0_ is an intercept of the linear model. Note that, in real analysis, some of 𝑥_j,𝑡_∗ may be time-delayed coordinates of an interacting variable. The linear model is fit to the other vectors in the state space, but points that are close to the target point, 𝑿_𝑡_∗, are given greater weighting (i.e., locally weighted linear regression). In the standard S-map, the distances between the target point, 𝑿_𝑡_∗, and other points are measured by Euclidean distance. However, Euclidean distance cannot be a good measure if the dimension of the state space is high. In the MDR S-map, the distance between the target point and other points is measured by the multiview distance (Chang et al., 2021), which can be calculated by ensembling distances measured in various low-dimensional state spaces (multiview embedding; see Ye & Sugihara, 2016). In addition, to reduce the possibility of overestimation and to improve forecasting skill, regularization (i.e., ridge regression) is also applied (Cenci et al., 2019). As in the UIC analysis, we included causal climate variables as conditional variables. In the present study, we implemented the MDR S-map in our custom R package, “macam” v0.0.10 (Ushio, 2022) and used it for computation.

### 2.9 Data and code availability

All scripts and raw data except for the sequence data used in the present study are available on Github (https://github.com/ong8181/ogura-rice2017) and archived on Zenodo (https://doi.org/10.5281/zenodo.17317363). High-throughput sequence data was deposited in DDBJ Sequence Read Archives (DRA) (BioProject ID = PRJDB9326, DRA Run ID = DRR709379-DRR709530 [16S rRNA], DRR709531-DRR709682 [18S rRNA], DRR709683-DRR709834 [COI], and DRR709835-DRR709986 [ITS]).

## 3. RESULTS

### 3.1 Rice growth trajectory and yield survey

During the monitoring period in 2017, the daily mean air temperatures were about 20°C in May, peaked at 30°C in August, and decreased to 20–25°C in September (Fig. 1b). In 2017, increased rainfall was observed due to seasonal frontal activity in June and typhoons in July, August, and September, which was negatively correlated with daily sunlight hours (Fig. 1c, d). In general, daily air temperatures during the monitoring period were similar or slightly higher than those during the normative year, while the rainfall was greater than during the normative year because of the seasonal frontal activity and typhoon activity.

The rice growth rate, SPAD, and the number of stems were consistently higher in the conventional paddy field than in the no-fertilizer paddy field, except for the first-differenced SPAD values (the original values; Fig. 2a–c: the first-differenced values; Fig. 2d–f) (*P* < 0.0001, Table S2). Large increases in the rice height and the number of stems were observed in late June and early July (Fig. 2a, c, d, f), while a large increase in SPAD value was observed in June (Fig. 2b, e). The total number of heads per rice individual and the total wet weight and total dry weight were also higher in the conventional paddy field than in the no-fertilizer paddy field (*P* < 0.0001; Fig. 2e, g and Table S3). These patterns were well reflected in the rice grain yields. The numbers of (total or fertile) grain counts, wet weight, and dry weight were higher in the conventional paddy field than in the no-fertilizer paddy field (*P* < 0.005; Fig. S2 and Table S3). Despite the lack of input of fertilizer, the no-fertilizer paddy field still produced (compared to the conventional paddy field production taken as 100%): total number of heads per 12 individuals: 56%, total grain count per head: 76%, and total grain dry weight per head: 77%, which corresponds to about 43% of total grain dry weight per individual (= 56% × 77%).

**Figure 2.**
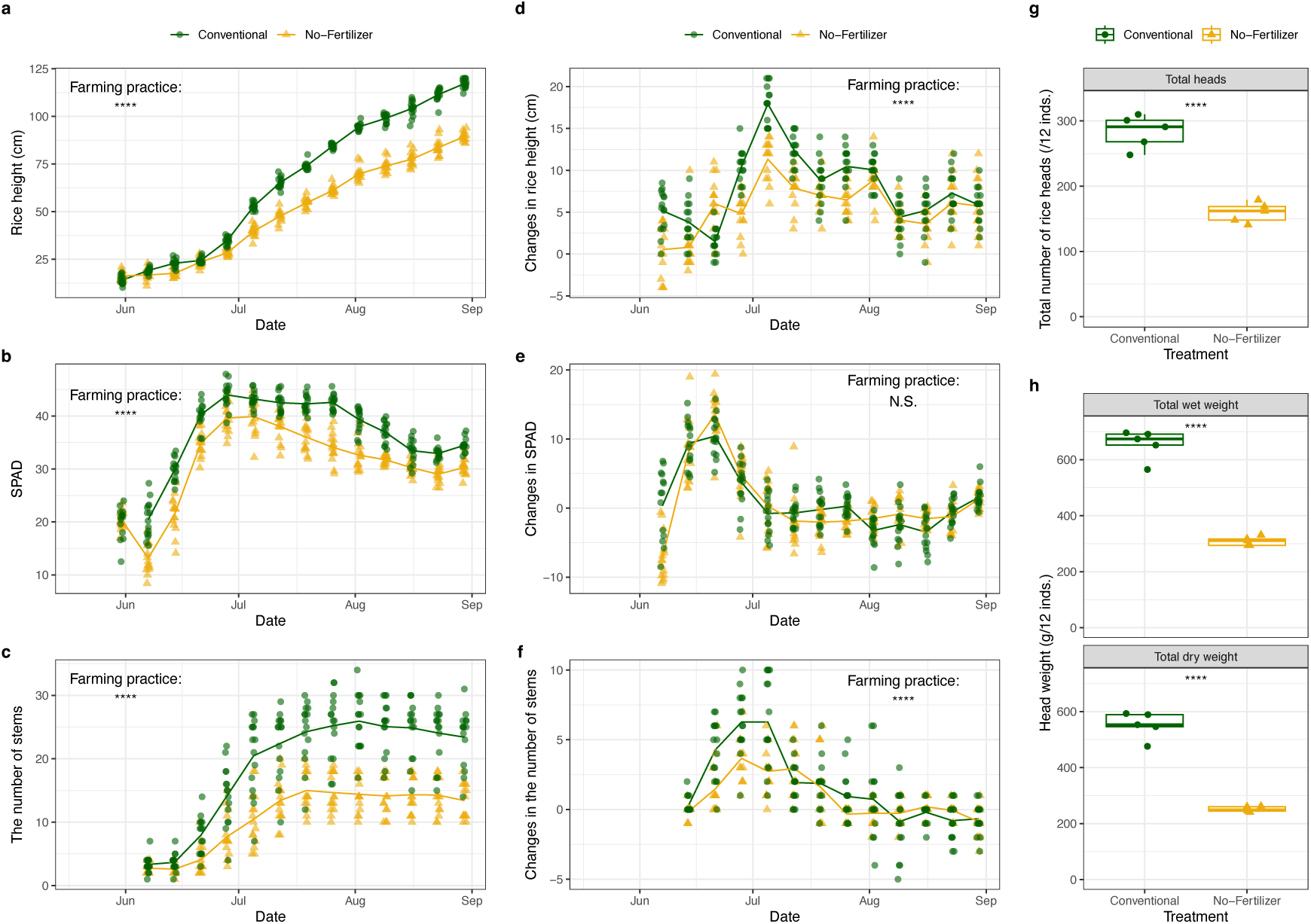
Rice growth patterns and yields. (**a**) Rice height (cm). (**b**) SPAD values. (**c**) The number of rice stems. (**d**) Changes in rice height per week. (**e**) Changes in SPAD values per week. (**f**) Changes in the number of rice stems per week. (**g**) The total number of rice heads per 12 rice individuals. (**h**) The total wet or dry weight of heads (g) per 12 rice individuals. Solid lines indicate mean values. Green color and and filled circle indicate data from the paddy fields using the conventional farming methods, and yellow color and filled triangle indicate data from the paddy fields using no fertilizers. \**P* < 0.05, \*\**P* < 0.01, \*\*\**P* < 0.001, and \*\*\*\**P* < 0.0001. “N.S.” indicates that the difference between the two farming practices is not statistically clear.

### 3.2 Plant specialized metabolites, insect abundance, and plant cover on the ridge

We detected eight dominant plant specialized metabolites: (*Z*)-3-hexenal, (*E*)-2-hexenal, (*Z*)-3-hexen-1-ol (leaf alcohol), 2-4-heptadienal, linalool, β-caryophyllene, methyl salicylate (MeSA), and diterpenoids (Fig. 3a). The concentrations of (*Z*)-3-hexenal and (*E*)-2-hexenal were higher than those of the other metabolites. The levels of (*Z*)-3-hexenal and β-caryophyllene were greater in the early period (June) compared to the late period, while 2-4-heptadienal showed higher concentrations in the late period. (*E*)-2-hexenal, (*Z*)-3-hexen-1-ol, and MeSA were detected throughout the monitoring period, although their temporal patterns were unclear. Linalool and diterpenoids were observed only sporadically. Overall, the differences between the conventional and no-fertilizer paddy fields were relatively unclear, but we identified statistically clear effects of the farming practice on (*Z*)-3-hexenal, (*E*)-2-hexenal, and methyl salicylate (Fig. 3a; Table S2).

**Figure 3.**
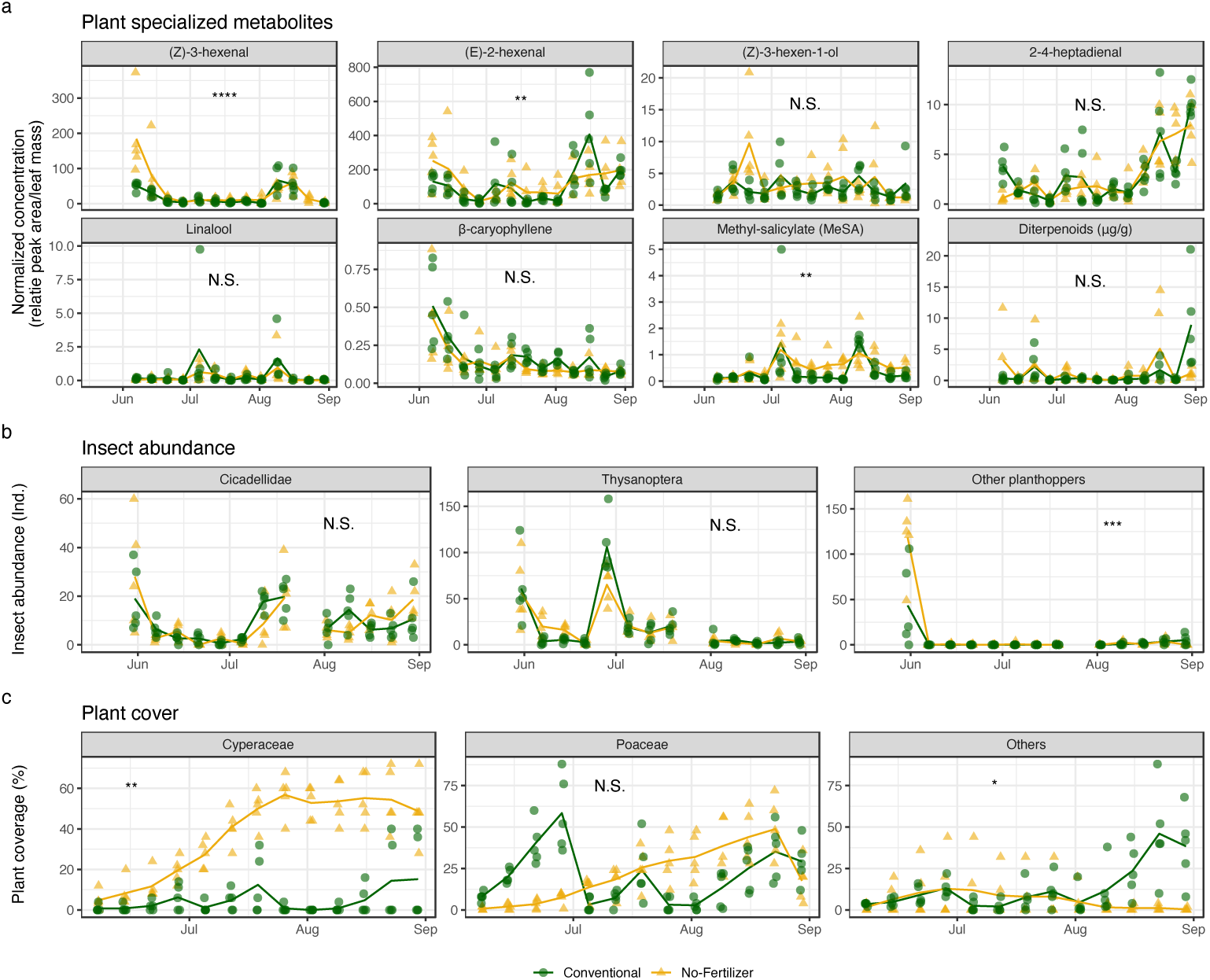
Plant specialized metabolites, insect abundance, and plant cover in the paddy fields. (**a**) Plant specialized metabolites. Each panel indicates a different type of plant specialized metabolites; (*Z*)-3-hexenal, (*E*)-2-hexenal, (*Z*)-3-hexen-1-ol, 2-4-heptadienal, linalool, 𝛽-caryophyllene, and methyl salicylate (MeSA) indicate plant volatile (relative peak area/fresh leaf weight), while diterpenoids (𝜇g/g fresh leaf weight) indicates phytoalexin. (**b**) Insect abundance trapped by the sticky plate method set on the ridge between paddy fields. Each panel shows a different insect group. (**c**) Plant coverage (%) on the ridge area between the paddy fields. Each panel shows a different plant type. Solid lines indicate mean values. Green color and and filled circle indicate data from the conventional paddy field, and yellow color and filled triangle indicate data from the no-fertilizer paddy field. \**P* < 0.05, \*\**P* < 0.01, \*\*\**P* < 0.001, and \*\*\*\**P* < 0.0001. “N.S.” indicates that the difference between the two farming practices is not statistically clear.

Thrips (Thysanoptera) were the most commonly observed insect taxon during the monitoring period (Fig. 3b), primarily in the first half of the monitoring (June to July). Leafhoppers (Cicadellidae) were also frequently observed, though in smaller numbers than thrips, except in late June. Other planthoppers were observed almost exclusively in early June. Statistically clear effects of the farming practice were detected only for other planthoppers (*P* < 0.001; Fig. 3b; Table S2).

A total of 14 plant species from six families were recorded on the paddy ridge (Figs. 3c and S3). Species from the Cyperaceae family were more dominant in the no-fertilizer paddy field (*P* < 0.01; Fig. 3c; Table S2). No statistically clear difference was found in Poacea between the two paddy fields. In contrast, other plant species were more dominant in the conventional paddy field, particularly in the later part of the monitoring period (*P* < 0.05; Fig. 3c; Table S2). In the conventional paddy field, an annual graminoid, *Digitaria ciliaris*, was the most dominant species in June. From July to August, an annual graminoid, *Dinebra chinensis*, and two annual forbs, *Euphorbia maculata* and *Portulaca oleracea*, became dominant. In the no-fertilizer rice field, a perennial forb, *Ixeris japonica*, dominated from June to early July, followed by two perennial sedges, *Fimbristylis diphylloides* and *Cyperus brevifolius*, which dominated in late July to August.

The overall patterns of plant cover and specialized metabolites were analyzed using NMDS (Fig. 4). Insect community compositions were not compared, as only three insect groups were examined in this study. No statistically clear differences were found in the composition of specialized metabolites between the two paddy fields (number of weeks, *P* = 0.0147; farming practice, *P* > 0.05; interaction, *P* > 0.05). However, a statistically clear difference was observed in plant cover composition between the conventional and no-fertilizer paddy fields (number of weeks, *P* < 0.0001; farming practice, *P* < 0.0001; interaction, *P* < 0.0001), which aligns with the patterns observed in the individual metabolites and plant species presented in Figs. 3 and S3.

**Figure 4.**
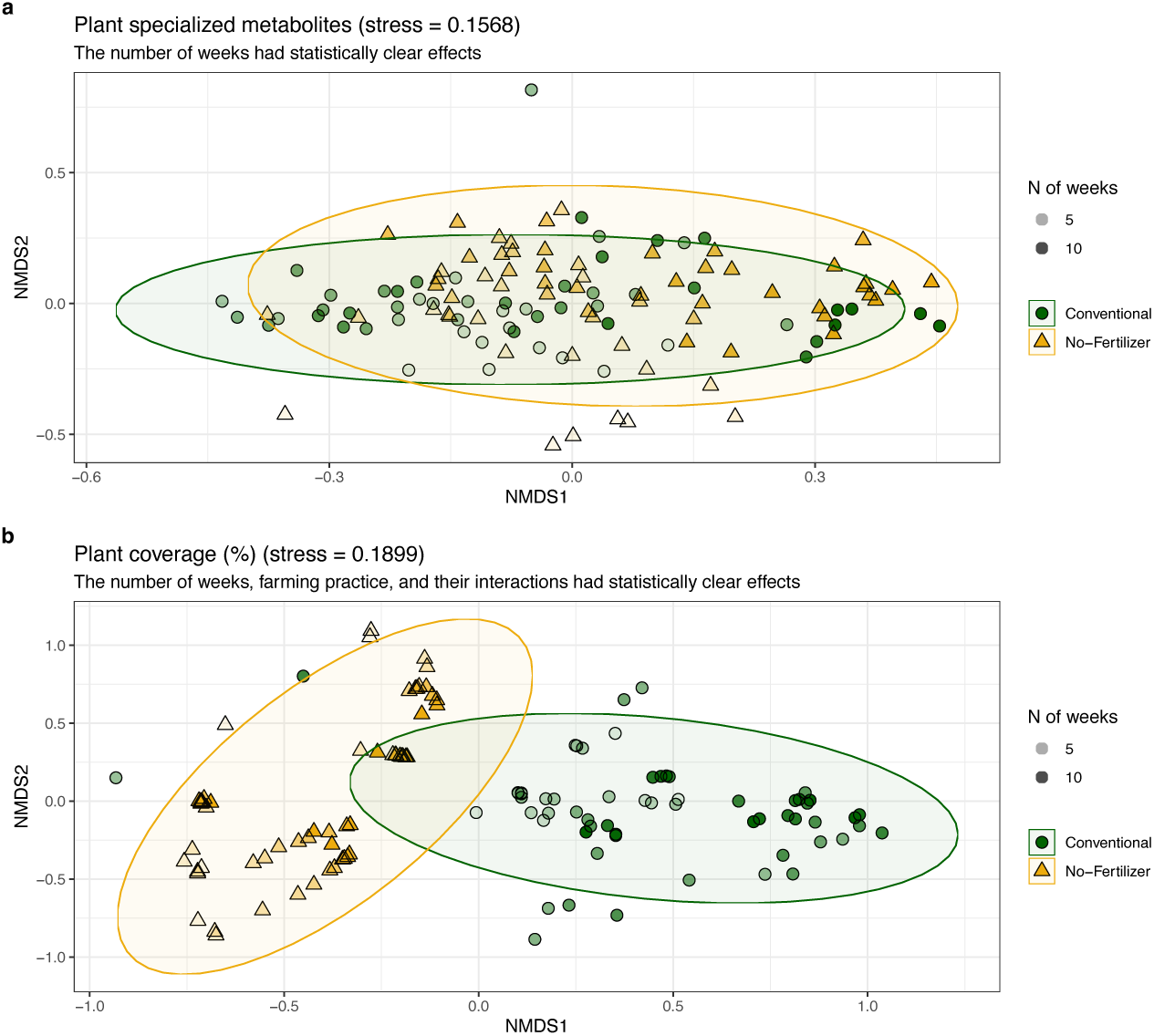
Non-metric dimensional scaling (NMDS) of plant communities on the paddy ridges and plant specialized metabolites. (**a**) NMDS of the plant communities on the paddy ridges based on Bray-Curtis dissimilarity (stress value = 0.1899). (**b**) NMDS of the plant specialized metabolites based on Bray-Curtis dissimilarity (stress value = 0.1568). The density of each color indicates the number of weeks since the rice planting in the rice fields. Green and and filled circle indicate data from the conventional paddy field, and yellow and filled triangle indicate data from the no-fertilizer paddy field.

### 3.3 Ecological community dynamics detected by quantitative eDNA metabarcoding

Quantitative eDNA metabarcoding was conducted using water samples from the paddy field. A total of 21,638,562 high-quality sequence reads (>Q30 = 94.17%) were obtained from four MiSeq runs (7,274,858 reads with >Q30 = 89.19% for 16S; 7,352,339 reads with >Q30 = 98.09% for 18S; 7,749,778 reads with >Q30 = 97.13% for COI; and 2,899,016 reads with >Q30 = 87.25% for ITS). Generally, we detected negligible non-target DNA sequence reads from negative controls (Fig. S4). While some negative controls detected non-target DNA sequences, these were mostly specific to the negative controls and did not affect subsequent analyses. Thus, we concluded that no serious contamination occurred during our sampling and library preparation processes. After DADA2 processing, 20,067,623 reads remained (Table S4). Following the conversion to copy numbers per ml of water and the filtering of very rare and infrequently detected OTUs, we identified 1,213 OTUs (Table S5). Table S6 provides a summary of eDNA copy numbers, sample metadata, and taxonomic information.

Regarding the temporal trend of total eDNA concentration, we observed contrasting patterns between the two paddy fields; total eDNA concentration decreased in the conventional paddy field while it increased in the no-fertilizer rice field during the monitoring period (Fig. 5a). A statistically clear effect of the farming practice on total eDNA concentration was found (*P* < 0.0001; Fig. 5a).

**Figure 5.**
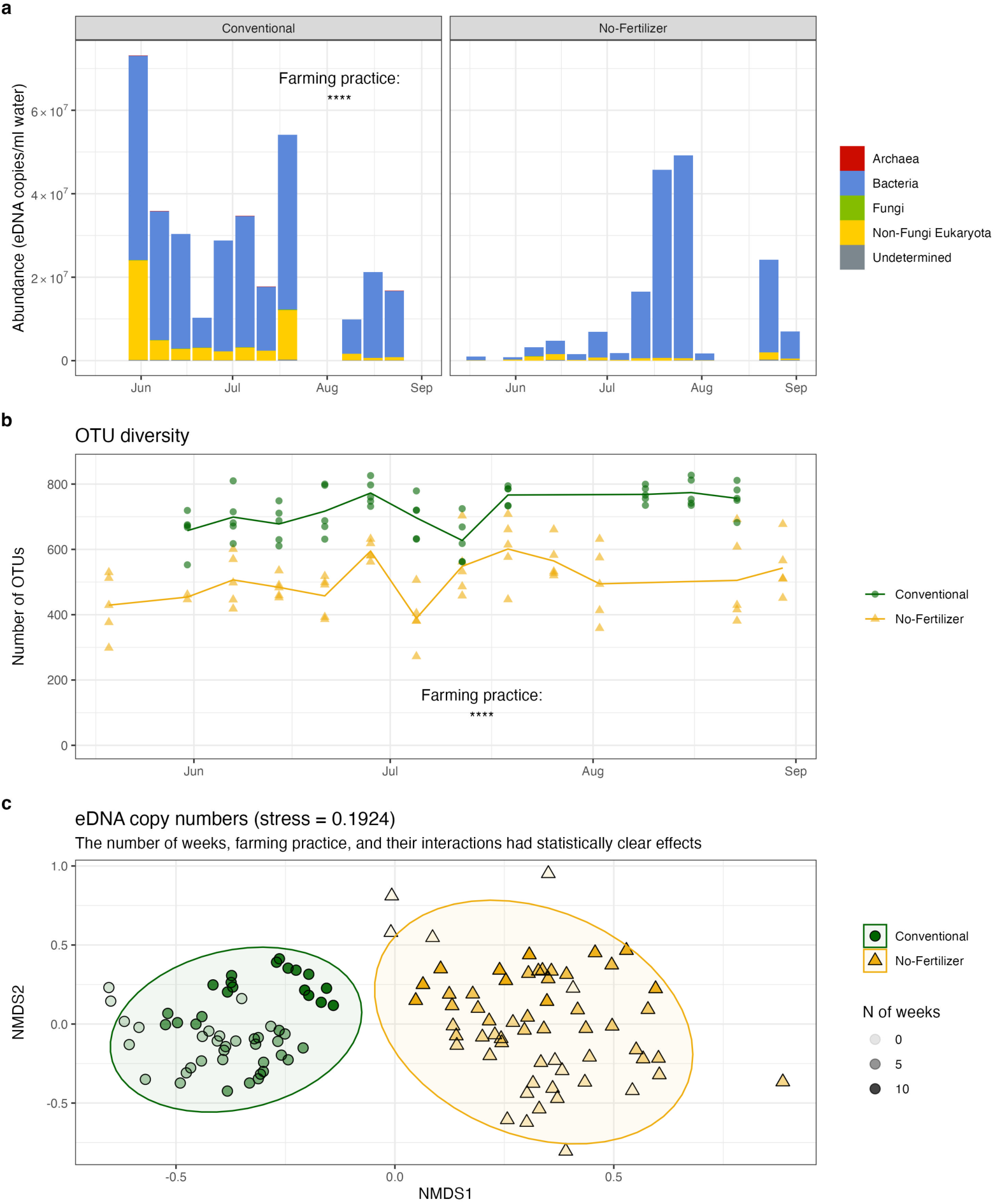
Environmental DNA in water samples collected from the two paddy fields. (**a**) The abundance (eDNA copies/ml water) of the ecological communities in the water samples. Each panel indicates the conventional or no-fertilizers paddy field. Colors indicate different taxonomic groups. (**b**) The number of OTUs detected in the water samples. Solid lines indicate mean values.(**c**) NMDS of the eDNA-based ecological community data based on Bray-Curtis dissimilarity (stress value = 0.1924). Green and and filled circle indicate data from the conventional paddy fields, and yellow and filled triangle indicate data from the no-fertilizer paddy field. \**P* < 0.05, \*\**P* < 0.01, \*\*\**P* < 0.001, and \*\*\*\**P* < 0.0001. “N.S.” indicates that the difference between the two farming practices is not statistically clear.

Bacteria were the most dominant taxa detected (Figs. 5a), with dominant genera including *Paenibacillus* sp., *Flavobacterium* sp., *Pedobacter* sp., and several species belonging to Actinomycetes and Comamonadaceae (Figs. S5, S6; Table S7). Most major bacterial and fungal phyla showed higher abundance in the conventional paddy field compared to the no-fertilizer paddy field (Fig. S6). However, Actinomycetota and Bacillota were more abundant in the no-fertilizer paddy field after July–August. Additionally, Fusobacteriota showed increased abundance during June–July in the no-fertilizer paddy field. The dominant non-bacterial taxa included green algae (Desmidiales), freshwater algae (*Gomphonema parvulum*), freshwater ostracods (Cyprididae), and small crustaceans (*Monia* sp.) (Table S7), with generally higher abundances in the conventional paddy field than in the no-fertilizer paddy field. Photosynthetic taxa, including Bacillariophyta, Chlorophyta, and Streptophyta, were also more abundant in the conventional paddy field compared to the no-fertilizer paddy field (Fig. S7).

The number of OTUs detected, which serves as an index of species richness, was consistently higher in the conventional paddy field than in the no-fertilizer paddy field (*P* < 0.0001; Fig. 5b; Table S2). In the conventional paddy field, the number of detected OTUs ranged from approximately 600 to 800, consistently exceeding that in the no-fertilizer paddy field.

The ecological community compositions identified through the quantitative eDNA metabarcoding were also analyzed using NMDS (Fig. 5c). There was a statistically clear difference between the two paddy fields, as well as a clear seasonal pattern (number of weeks, *P* < 0.0001; farming practice, *P* < 0.0001; interaction, *P* < 0.0001). In the conventional paddy field, the ecological community compositions were characterized by species belonging to Actinomycetes, γ-Proteobacteria, β-Proteobacteria, Verrucomicrobia, Oligoflexia, and Sphingobacteriia (Table S8; Fig. S8). The abundances of most of these species were higher during the first half of the monitoring period in the conventional paddy field, with differences between the two paddy fields diminishing after August. In the no-fertilizer paddy field, the ecological community composition was predominantly characterized by the dominance of Actinomycetes (PRO_Taxa00038; Table S8; Fig. S8). The abundance of PRO_Taxa00038 was higher in the no-fertilizer paddy field than in the conventional paddy field until August.

### 3.4 Effects of ecological variables on the rice performance

Based on the time series data, we detected causal variables that affect rice performance, specifically height growth (Fig. 6 and Table S9) (For results on SPAD values and stem counts, see Figs. S9, S10 and Tables S10, S11). Using UIC, we detected 16 ecological variables that causally impacted rice growth. The OTUs with the strongest positive effects on rice height growth were Undetermined taxon (ITS_Taxa00138), β-Proteobacteria (PRO_Taxa00019), and Bacteria (PRO_Taxa00332) (Fig. 6a). These species were more abundant in the conventional rice field (Table S9 and Fig. S11), contributing to improved rice growth in that environment. While our analysis included both microbes and macrobes (e.g., invertebrate taxa), only *Digitaria ciliaris* (Poaceae) on the ridge had statistically clear causal effects on rice growth. Several bacterial species from Bacteroidota, Pseudomonadota, Burkholderiales, *Flavobacterium* sp., and *Polynucleobacter cosmopolitanus* also showed clear causal influences on rice growth. Additionally, a species from Chlamidomonadales affected rice growth. Species from Cochliopodium, Chlorophyceae, Ploima, Actinomycetes, eukaryota, and bacteria has statistically clear effects on SPAD values (Fig. S9), while only two species from Silvanigrellaceae and Chlorophyceae had statistically clear effects on stem counts (Fig. S10).

**Figure 6.**
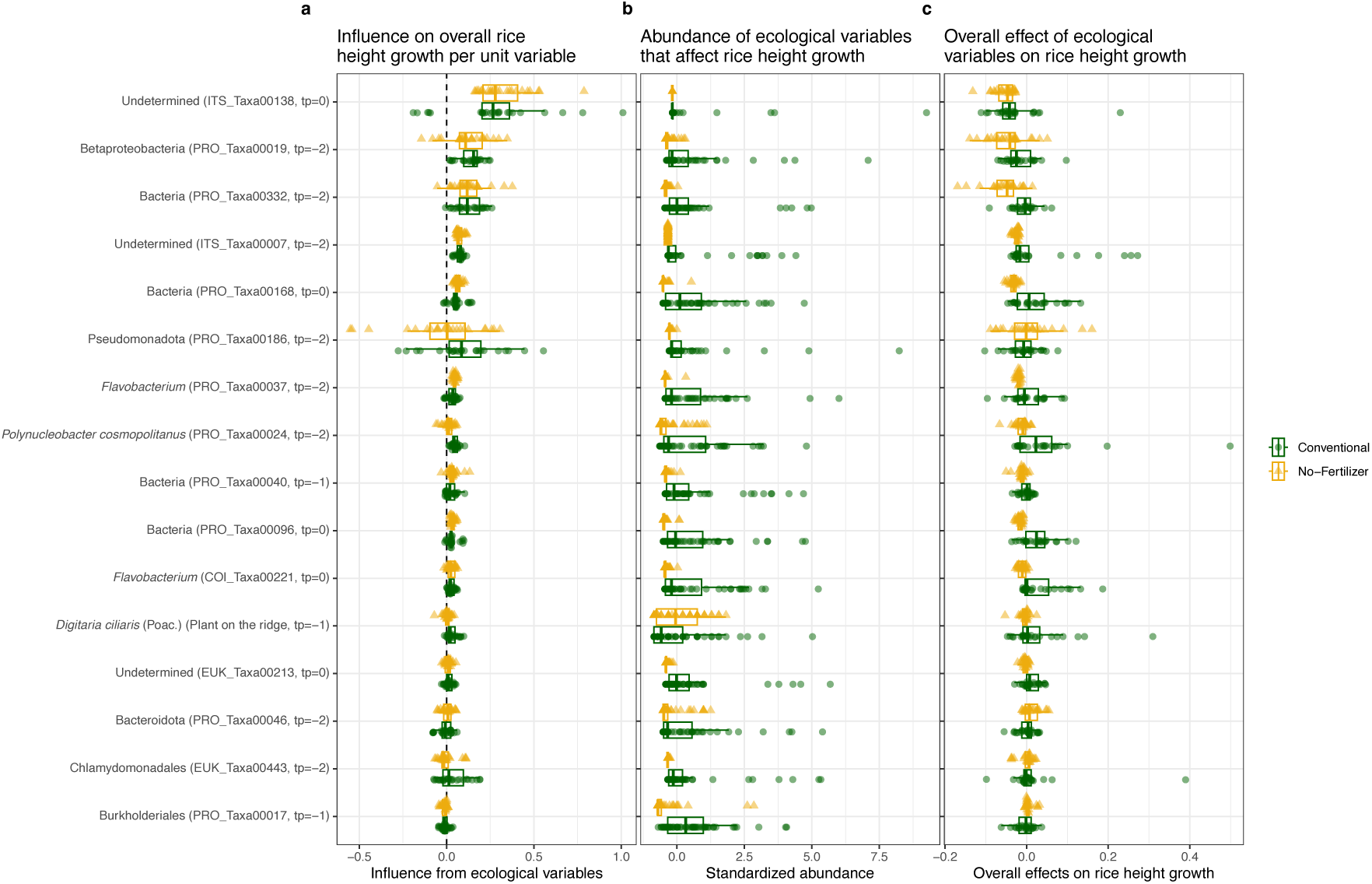
Effects of ecological variables on rice height growth in the conventional and no-fertilizer paddy fields. Each panel shows (**a**) the effects of ecological variables on overall rice height growth per unit variable (i.e., coefficients of the MDR S-map), (**b**) the abundance of ecological variables, and (**c**) the overall effects of ecological variables on rice height growth (calculated by “the MDR S-map coefficient × Abundance”), respectively. Green and and filled circle indicate data from the conventional paddy fields, and yellow and filled triangle indicate data from the no-fertilizer paddy fields. The 𝑦-axis labels include assigned taxa names, taxa code, and effect delay (𝑡𝑝; e.g., 𝑡𝑝 = −1 indicates the taxon has effects on the rice performance one week later).

Importantly, the impact of these ecological members per abundance on overall rice performance was largely consistent between the two paddy fields (Figs. 6a, S9a, and S10a and Tables S9–S11). Differences in their overall contributions to rice growth between the two fields (Figs. 6c, S9c, and S10c and Tables S9–S11) were primarily attributed to variations in the abundance of these causal species (Figs. 6b, S9b, and S10b and Tables S9–S11). Most causally influential species were more abundant in the conventional paddy field compared to the no-fertilizer paddy field (Fig. S11). The abundances of ecological community members positively influencing rice growth were generally higher in the conventional rice field than in the no-fertilizer rice field.

## 4. DISCUSSION

In the present study, we conducted thorough monitoring to identify ecological variables that affect rice growth performance in the two paddy fields with different farming practices. We observed clear differences in rice performance regarding height growth, SPAD values, stem counts, and yields. Despite the lack of fertilizer inputs, the no-fertilizer paddy field still produced approximately 40–50% of the rice yields compared to the conventional paddy field. The rice yield in the no-fertilizer field was lower than that reported in previous research (e.g., Okumura, 2002), likely due to differences in rice variety (*Hinohikari* in the present study versus *Beniasahi* in the previous research). Ecological variables, including plant species on the paddy ridge and ecological communities identified using the quantitative eDNA metabarcoding, also showed clear differences between the fields. Using time series analysis, we identified ecological variables that influence rice performance.

### 4.1 Contrasting ecological communities in the two paddy fields

First, we observed statistically clear differences in ecological communities between the two paddy fields. The conventional paddy field exhibited a lower abundance of Cyperaceae species on the ridge (Fig. 3), a higher total eDNA concentration in June and July (Fig. 5a), and greater OTU richness (Fig. 5b). Specific OTUs from γ-proteobacteria, Chlorophyceae, Verrucomicrobiota, Actinomycetes, and other taxa displayed distinct differences between the two fields (Fig. S8 and Table S7). Given that the climate variables were nearly identical in the two paddy fields, the farming practices, namely the use of chemical fertilizers and pesticides, were the primary factors driving these differences.

#### 4.1.1 Plant specialized metabolites and insect abundance

We observed only minor differences in plant specialized metabolites and insect abundance between the two paddy fields (Fig. 3). Among the metabolites measured, (*Z*)-3-hexenal, (*E*)-2-hexenal, and methyl salicylate showed slightly higher concentrations in the rice leaves of the no-fertilizer paddy field compared to those of the conventional field (Fig. 3a and Table S2). (*Z*)-3-hexenal and (*E*)-2-hexenal are common green leaf volatiles (GLVs) that can be released in response to mechanical damage and can activate defense genes against certain pathogenic fungi (Matsui & Koeduka, 2016). Methyl salicylate is a type of insect-induced plant volatile and can be emitted by rice seedlings in response to insect herbivory (Zhao et al., 2010). Since the no-fertilizer paddy field did not receive any pesticides, rice plants there may have been more frequently affected by arthropodal pests and microbial pathogens. The abundance of an OTU belonging to Actinomycetes (PRO_Taxa00038) was approximately 100 times higher in the no-fertilizer paddy field (Fig. S8), which may have contributed to the increased emission of the two GLVs.

The abundances of the three insect groups were generally similar (Figs. 3, 4), but a statistically clear difference was found in the category of “other planthoppers,” likely due to their higher abundance in the no-fertilizer paddy field during the first monitoring event. Interestingly, the increased abundance of “other planthoppers” in early June coincided with higher concentrations of (*Z*)-3-hexenal and (*E*)-2-hexenal, suggesting that herbivory damage may have contributed to the elevated levels of GLVs in the no-fertilizer paddy field. Overall, however, the differences in plant specialized metabolites and insect groups were subtle, as indicated by the NMDS (Fig. 4a). This suggests that fertilizers and pesticides have weaker impact on plant specialized metabolites and insect groups compared to their effects on plant communities on the paddy ridge and other ecological communities (Figs. 4b and 5), at least in the two paddy fields studied. The weaker effects of pesticides on the insect communities may also have resulted from the fact that our survey plots were located within 50–70 m of each other, so insects could migrate between the conventional paddy field and the no-fertilizer paddy field (Fig. S1a).

#### 4.1.2 Plant communities on the paddy ridge

In the conventional paddy field, all dominant species were annuals, while perennials were more prevalent in the no-fertilizer paddy field. The plant species observed on the paddy ridges are species commonly found in agricultural fields across Japan. Pesticide application for weed control eliminates existing plants, allowing fast-growing annuals to dominate. In contrast, mowing does not disrupt the underground structures, allowing perennials with low growing points or those that reproduce vegetatively via rhizomes or stolons to thrive. In particular, two perennial species from the Cyperaceae family, *Cyperus brevifolia* and *Fimbristylis diphylloides*, were particularly dominant in the no-fertilizer paddy field. Many Cyperaceae species are well adapted to open, sunny, moist environments, which constitute major components of paddy ecosystems (Bryson & Carter, 2008). Specifically, *Cyperus brevifolia* develops a robust rhizome network, enabling it to quickly establish a dense, space-occupying mat (Shaw et al., 2011).

#### 4.1.3 Ecological community compositions detected by eDNA metabarcoding

The increased total eDNA concentration in the conventional paddy field, primarily due to higher microbial abundance, was likely influenced by fertilizer inputs. While nitrogen addition to the soil can lead to a reduction in microbial biomass (Treseder, 2008), it can also enhance phytoplankton biomass in freshwater ecosystems (Søndergaard et al., 2017). In the present study, major photosynthetic taxa, including Chlorophyta, were more abundant in the conventional paddy field (Fig. S7), likely due to the higher nitrogen input. The greater abundance of these photosynthetic taxa contributes to increased carbon input into the system, which may boost microbial biomass (and total eDNA concentration) in the conventional paddy field. Additionally, the higher productivity of rice further adds to the carbon input. Indeed, Kamata et al. (1991) reported a greater concentration of soil carbon in conventional paddy soil compared to no-fertilizer paddy soil (in the no-fertilizer paddy soil studied in Ritto, Shiga Prefecture, Japan).

The nitrogen fertilizer and increased carbon input were likely the primary factors driving the differences in community composition (Figs. 5 and S6–S8). Changes in community composition in response to nutrient addition or carbon input have frequently been reported in paddy fields (e.g., Dong et al., 2014; Z. Zhao et al., 2019). Actinomycetes (PRO_Taxa00038), which contributed to the differences between the two paddy fields, share the same 16S sequence fragment with sequences detected in Neotropical floodplain lakes (Targueta et al., 2023), suggesting a potential association with nutrient-poor conditions typical of natural ecosystems rather than agricultural environments. Several Chlorophyceae OTUs (EUK_Taxa00281, EUK_Taxa00423, and EUK_Taxa00170) were more abundant in the conventional paddy fields than in the no-fertilizer fields, and Chlorophyceae have been detected in rice paddy fields where light and nutrient availability are high (Sahu, 2012).

### 4.2 Influences of ecological communities on the rice performance

Our time series analysis detected ecological variables that can causally influence rice performance (Figs. 6, S9–S11 and Tables S9–S11). Among the causal taxa affecting rice height growth, some, such as ITS_Taxa00138 and ITS_Taxa0007, showed much higher abundances in the conventional paddy field than in the no-fertilizer field. Although we could not assign species names based on the OTU sequences, a common characteristic of these taxa was their dominance during the early monitoring period, particularly in June (Fig. S11). This pattern also applied to the only causal plant species on the paddy ridge, *Digitaria ciliaris* (Fig. S3). While we could not identify the genus or species for most taxa, exceptions included *Flavobacterium* spp. (PRO_Taxa00037 and COI_Taxa00221) and *Polynucleobacter cosmopolitanus* (PRO_Taxa00024), which were also dominant in the conventional paddy field in June. *Flavobacterium* is a genus of Gram-negative bacteria found in various ecosystems, and *F. keumense*, a species related to COI_Taxa00221, was isolated from freshwater ecosystems in South Korea (Ekwe et al., 2017). *Polynucleobacter cosmopolitanus* is a free-living, cosmopolitan freshwater bacterioplankton frequently detected in ecosystems in Japan (Watanabe et al., 2012). Physiological and ecological information about these species is limited, making it difficult to infer the exact mechanisms by which these taxa influence rice height growth in this study. Future research could leverage the spatio-temporal data on these beneficial taxa, i.e., their general dominance in June and higher detection rates in conventional paddy fields, to focus on identifying their ecological and physiological roles.

Similarly, while we detected statistically clear causal effects of several taxa (e.g., Cochliopodium, Ploima, Actinomycetes, Silvanigrellaceae, and Chlorophyceae) on SPAD values and the number of stems (Figs. S9, S10), it remains difficult to determine the exact mechanisms by which they influence rice performance. Interestingly, some of these species showed increased abundance during the middle of the monitoring period (from late June to July). Thus, the mechanisms through which these taxa affect rice performance may differ from those influencing rice height growth. Focusing on taxa that demonstrate clear temporal patterns, such as PRO_Taxa00050 (Bacteria) and COI_Taxa00134 (Ploima, a rotifer species) (Fig. S11), could provide insights into their roles in the paddy fields.

### 4.3 Potential factors that contribute to the maintenance of the rice performance in the paddy fields

While we could not determine the exact mechanisms by which ecological communities influence rice performance in the paddy fields, our analysis provide insights into the differences between the two fields. Importantly, the per-abundance effects of key ecological community members on overall rice performance were generally similar across both paddy fields (Figs. 6a, S9a–S10a). Statistically clear differences in the effects between the farming practices were observed for only a few taxa (Tables S9–S11; three out of 16 species, zero out of seven species, and one out of two species). In contrast, 13 out of 16 species, four out of seven species, and both species in one category showed statistically clear differences in abundance between the two paddy fields, with most showing lower abundance in the no-fertilizer field than in the conventional field (Fig. S11). These findings suggest that, despite the differences in rice yields, growth trajectories, and ecological communities between the two paddy fields, the interactions between rice and ecological communities were largely consistent. The abundance of influential species appears to contribute to the differences in rice growth performance and yields. While it is clear that nutrient inputs (i.e., N-P-K fertilizers) enhanced rice growth and yield in the conventional paddy fields, the abundance and composition of ecological community members may also support rice growth. This indicates that introducing these species could potentially improve rice growth and yields in the no-fertilizer paddy field.

### 4.4 Advantages, limitations, implications and future direction

In this study, we integrated monitoring of rice performance, collection of plant specialized metabolites, plant cover surveys on the ridge, insect survey, eDNA-based ecological community monitoring, and a nonlinear time series causality detection to understand the impact of ecological variables in actual rice paddy fields. The integration of field-based monitoring with advanced causality detection methods enabled the detection of potential causal ecological variables influencing rice growth in the real paddy fields.

One advantage of our research framework is its ability to elucidate ecological phenomena and complex causal interactions under field conditions. Manipulative experiments are among the most effective methods for understanding causal interactions in biological systems. However, while laboratory experiments can be conducted (provided that certain causal microbial species can be isolated and cultivated, or that other macrobial species can be easily maintained in the laboratory), the interactions observed in controlled settings may not necessarily reflect those in natural environments, as ecological interactions and resulting dynamics are often context-dependent and nonlinear (Hsieh et al., 2005; Sugihara et al., 2012; Ushio, 2022; Ushio, Hsieh, et al., 2018). In addition, multiple confounding factors, such as climate variables, exist in the fields, making the interpretation of manipulative experimental results difficult. Under these conditions, time-series analysis methods are some of the promising ways to understand complex interactions among variables. Indeed, time-series analysis-based causality detection has rapidly developed, and it has been applied especially when detailed manipulative experiments are not feasible, as seen in atmospheric sciences, neuroscience, and environmental sciences (Runge et al., 2019; Sugihara et al., 2012; Suzuki et al., 2022; Tajima et al., 2017). UIC enables robust causality detection based on clear mathematical definitions, avoiding pseudo-correlations (Osada et al., 2023), while the MDR S-map quantifies interaction strength even when the number of causal variables exceeds the number of observations, circumventing the curse of dimensionality (Chang et al., 2021). Therefore, although their application in agricultural sciences is still limited, integrating field-based monitoring with advanced time-series analysis can provide a more realistic understanding of agricultural systems.

Nonetheless, we must address the technical limitations of our study for future research. First, while quantitative eDNA metabarcoding offers a comprehensive and quantitative ecological time series (Ushio, 2022; Ushio, Murakami, et al., 2018), several challenges remain, particularly regarding abundance estimation and species identification. The relationship between eDNA concentrations and species abundance, especially for macro-organisms, remains an open question (Harrison et al., 2021). eDNA concentrations should be considered “an index” of species abundance, as the rates of eDNA release and degradation vary by species and environmental conditions (Dejean et al., 2011). For microbial species, variations in the copy numbers of target genes, particularly 18S rRNA (Godhe et al., 2008), complicate the estimation of abundance or biomass from metabarcoding data. Additionally, assigning species names using short-read amplicon data is often challenging. Long-read sequencing-based eDNA metabarcoding or shotgun metagenomic analysis may address these issues associated with short-read amplicon sequencing, as they can provide more detailed information, including intra-specific and individual variations within ecological communities (Baloğlu et al., 2021; Nousias et al., 2025).

Another limitation is the number of time points in our time series. Although we made substantial efforts to measure multiple ecological variables in the two paddy fields, our monitoring was conducted weekly. While we detected “statistically clear” causal effects of ecological variables, our sampling design did not allow us to capture dynamics occurring within a timeframe shorter than one week. For instance, rice gene expression dynamics can vary across multiple time scales, including seasonal, daily, hourly, and even shorter intervals (Nagano et al., 2012). Utilizing more frequent and high-throughput field monitoring technologies, such as unmanned aerial vehicles (UAVs) for rice monitoring (Chen et al., 2024) and high-throughput, low-cost RNA-seq methods (Kamitani et al., 2019), would greatly enhance the efficiency and accuracy of detecting causal ecological variables in field conditions.

### 4.5 Conclusions

In the present study, we integrated various ecological monitoring methods and a time series-based causality detection approach to explore the role of ecological variables in sustaining rice performance in both conventional and no-fertilizer paddy fields in Japan. Our time series analysis revealed several previously unrecognized ecological community members that causally influenced rice performance. Interestingly, these members showed similar per-abundance effects on rice in both fields but their abundance differed, suggesting that similar mechanisms underlie the differing rice growth patterns, with abundance potentially driving some of the differences observed. While we could not fully elucidate the detailed mechanisms by which these communities affect rice performance, we demonstrated that our framework—integrating field-based monitoring with advanced time series analysis— facilitates an understanding of the role of ecological communities in rice paddy fields. Future studies that incorporate more advanced monitoring and data analysis techniques will enhance our understanding and management of rice paddy fields, promoting more sustainable crop production.

## Acknowledgments

We thank Itsuo Kitagawa, Keisuke Katsura, and Satomi Yoshinami for assistance in the field survey, Asako Kawai for assistance in the DNA library preparations, and Akifumi Sugiyama for assistance in the field sampling. This research was supported by KAKENHI (B) 17H03811 and 20H02922 (to KO), PRESTO (JPMJPR16O2) from the Japan Science and Technology Agency (JST), the Hakubi Project in Kyoto University (to MU), and The Hong Kong University of Science and Technology Startup Fund (to MU).

## Author Contributions

MU, HS, RO, KS, and JT conceived and designed research; MU, HS, YS, KO, RO and KS performed field monitoring and analyzed samples; MU performed time-series analysis; MU summarized and visualized the data; MU wrote the first draft; all authors discussed the results and completed the manuscript.

