## Supplementary Text for "Contrasting ecological communities in rice paddy fields under conventional and no-fertilizer farming practices"

#### Contents:

**Supplementary text**
**Supplementary methods**
**Figure S1.** Supplementary information for the rice field survey in 2017
**Figure S2.** Rice grain data
**Figure S3.** The coverage (%) of plant species on the paddy ridges
**Figure S4.** Sequence reads summary of the quantitative eDNA metabarcoding
**Figure S5.** Top 12 most-abundant taxa detected by the quantitative eDNA metabarcoding
**Figure S6.** Temporal dynamics of major bacterial and fungal phyla detected by the quantitative eDNA metabarcoding
**Figure S7.** Temporal dynamics of major non-fungal eukaryota detected by the quantitative eDNA metabarcoding
**Figure S8.** eDNA taxa that differentiate the conventional and no-fertilizer paddy fields
**Figure S9.** Influences of ecological variables on the SPAD values in the conventional and no-fertilizer paddy fields
**Figure S10.** Influences of ecological variables on the number of rice stems in the conventional and no-fertilizer paddy fields
**Figure S11.** Temporal dynamics of taxa that causally influenced rice performance
**Table S1.** Sample metadata
**Table S2.** Results of generalized additive mixed model (GAMM) of ecological variables
**Table S3.** Results of ANOVA for rice yield data
**Table S4.** eDNA sequence read summary
**Table S5.** Assigned taxa for the detected OTUs
**Table S6.** OTU table (eDNA copies numbers / ml water)
**Table S7.** List of most dominant taxa
**Table S8.** Top 20 taxa that contributed to the difference between the two paddy fields
**Table S9.** Results of linear mixed model (LMM) for the MDR S-map results of rice heights
**Table S10.** Results of linear mixed model (LMM) for the MDR S-map results of SPAD
**Table S11.** Results of linear mixed model (LMM) for the MDR S-map results for the number of rice stems

### Supplementary text

##### *The history and typical farming practice of the no-fertilizer paddy field*

The no-fertilizer paddy field, originally located in Ritto, Shiga Prefecture, Japan, covered approximately 1500 $m^2$ and consisted of two plots: 600 $m^2$ (Plot A) and 900 $m^2$ (Plot B). Irrigation water was sourced from a canal, entering Plot B and flowing through Plot A before draining into the outlet. Farming without fertilizers or pesticides was initiated by the rice field owner in 1951. In the no-fertilizer farming practice, all rice straw, husks, and manually removed weeds were transported off the field, leaving only a small amount of organic matter within the field, consisting of stubble, roots, and minimal weeds.

When the paddy field was in Ritto, Shiga Prefecture, the paddy field was prepared in spring, and seedlings were planted that had been grown without fertilizer. Initially, the planting was done by hand, but later, machine transplanting with pot-grown seedlings was employed. At first, four rounds of tilling were conducted every seven days starting about 20 days after planting, along with one round of manual weeding. Subsequently, from about 15 days after planting, three rounds of manual weeding and one round of mechanical weeding were carried out every ten days. No mid-season drainage was performed, and irrigation water was supplied until maturity. The cultivated variety was *Beniasahi* from the beginning, which is a long-stemmed, large-panicle, mid-late maturing variety, with seeds harvested from this rice field. The above information is available in Japanese on https://muhiken.or.jp/ritto/ (accessed as of 11 October 2025).

In 2006, after farming in Ritto ceased, the no-fertilizer paddy soil was moved to Uji, Kyoto, where no-fertilizer rice farming has continued. The farming management records for 2016 are as follows:

January 27 – Winter plowing
May 2 – Spring plowing
May 14 – Start of flooding
May 17 – First puddling (rough leveling)
May 25 – Second puddling (final leveling)
May 29 – Transplanting of rice seedlings (cultivar: *Beniasahi*)
June 8 – Manual weeding
June 22 – Manual weeding
September 27 – Drainage of the field
October 1 – Installation of drainage furrows using a ditching machine
October 14 – Rice harvesting

### Supplementary methods

##### *DNA extraction*

DNA was extracted using a DNeasy Blood & Tissue kit following a protocol described in a previous study (Ushio, 2019, 2022). First, the 2 ml of RNAlater solution in each filter cartridge were removed from the outlet under vacuum using the QIAvac system (Qiagen, Hilden, Germany), followed by a further wash using 1 ml of MilliQ water. The MilliQ water was also removed from the outlet using the QIAvac. Then, Proteinase K solution (20 µl), PBS (220 µl) and buffer AL (200 µl) were mixed, and 440 µl of the mixture was added to each filter cartridge. The materials on the cartridge filters were subjected to cell lysis by incubating the filters on a rotary shaker (15 rpm; DNA oven HI380R, Kurabo, Osaka, Japan) at 56˚C for 10 min. After cell lysis, filter cartridges were vigorously shaken (with zirconia beads inside the filter cartridges for 0.22-µm cartridge filters) for 180 sec (3200 rpm; VM-96A, AS ONE, Osaka, Japan). The bead-beating process was omitted for 0.45-µm cartridge filters, as the targets of the 0.45-µm cartridge filters were mainly macro-organisms' DNAs and we assumed that they were more vulnerable than those of microbial cells. The incubated and lysed mixture was transferred into a new 2-ml tube from the inlet (not the outlet) of the filter cartridge by centrifugation (3,500 g for 1 min). Zirconia beads in the 0.22-µm cartridge filters were removed by collecting the supernatant of the incubated mixture after the centrifugation. The collected DNA was purified using a DNeasy Blood & Tissue kit following the manufacturer's protocol. After the purification, DNA was eluted using 100 µl of the supplied elution buffer. Eluted DNA samples were stored at −20°C until further processing.

***Library preparation for the quantitative eDNA metabarcoding***

The first-round PCR (first PCR) was carried out with a 12-µl reaction volume containing 6.0 µl of 2 × KAPA HiFi HotStart ReadyMix (KAPA Biosystems, Wilmington, WA, USA), 0.7 µl of forward primer, 0.7 µl of reverse primer (each primer at 5 µM used in the reaction; with adaptor and six random bases), 2.6 µl of MilliQ water, 1.0 µl of DNA template and 1.0 µl of internal standard DNA mixture. We used the taxa-specific 1st PCR primer sets as follows: 515F-806R (Bates et al., 2011; Caporaso et al., 2011), Euk_1391f-EukBr (Amaral-Zettler et al., 2009), ITS1_F_KYO1-ITS_KYO2 (Toju et al., 2012), mlCOIintF (Leray et al., 2013) and HCO2198 (Folmer et al., 1994). Thermal cycle profile for each primer set was as follows:

*Prokaryote 16S rRNA (515F-806R)*

The thermal cycle profile after an initial 3 min denaturation at 95°C was as follows (35 cycles): denaturation at 98°C for 20 s; annealing at 60°C for 15 s; and extension at 72°C for 30 s, with a final extension at the same temperature for 5 min.

*Eukaryote 18S rRNA (Euk_1391f-EukBr)*

The thermal cycle profile after an initial 3 min denaturation at 95°C was as follows (35 cycles): denaturation at 98°C for 20 s; annealing at 62°C for 15 s; and extension at 72°C for 30 s, with a final extension at the same temperature for 5 min.

*Fungal ITS (ITS1_F_KYO1-ITS_KYO2)*

The thermal cycle profile after an initial 3 min denaturation at 95°C was as follows (35 cycles): denaturation at 98°C for 20 s; annealing at 55°C for 15 s; and extension at 72°C for

30 s, with a final extension at the same temperature for 5 min.

*Animal COI (mlCOIintF-HCO2198)*
The thermal cycle profile after an initial 3 min denaturation at 95°C was as follows (35 cycles): denaturation at 98°C for 20 s; annealing at 55°C for 15 s; and extension at 72°C for 30 s, with a final extension at the same temperature for 5 min.

Sequences for the primers are provided in the following section. Triplicate first PCRs were performed, and these replicate products were pooled in order to mitigate the PCR dropouts. The pooled first PCR products were purified using AMPure XP (PCR product: AMPure XP beads = 1:0.8; Beckman Coulter, Brea, California, USA). The pooled, purified, and 10-fold diluted first PCR products were used as templates for the second-round PCR.

The second-round PCR (second PCR) was carried out with a 24-μl reaction volume containing 12 μl of 2 × KAPA HiFi HotStart ReadyMix, 1.4 μl of each primer (each primer at 5 μM in the reaction volume), 7.2 μl of MilliQ water and 2.0 μl of template. Different combinations of forward and reverse indices were used for different templates (samples) for massively parallel sequencing with MiSeq. The thermal cycle profile after an initial 3 min denaturation at 95°C was as follows (12 cycles): denaturation at 98°C for 20 s; annealing at 68°C for 15 s; and extension at 72°C for 15 s, with a final extension at 72°C for 5 min. Twenty microliters of the indexed second PCR products were mixed, and the combined library was again purified using AMPure XP (PCR product: AMPure XP beads = 1:0.8).

Target-sized DNA of the purified library (ca. 440 bp for prokaryote 16S rRNA; ca. 320 bp for eukaryote 18S rRNA; ca. 470-600 bp for fungal ITS; ca. 510 bp for animal COI) was excised using E-Gel SizeSelect (ThermoFisher Scientific, Waltham, MA, USA) (Note that “the target size” includes Illumina P5/P7 adapter, Rd1SP/Rd2SP sequencing primer and sample-specific index sequences of which total length is 137 bp). The double-stranded DNA concentration of the library was quantified using a Qubit dsDNA HS assay kit and a Qubit fluorometer (ThermoFisher Scientific, Waltham, MA, USA). The double-stranded DNA concentration of the library was then adjusted using MilliQ water and the DNA was applied to the MiSeq (Illumina, San Diego, CA, USA). The prokaryote 16S rRNA, eukaryote 18S rRNA, fungal ITS and animal COI libraries were sequenced using MiSeq Reagent Kit V2 for 2 × 250 bp PE, MiSeq Reagent Kit V2 for 2 × 150 bp PE, MiSeq Reagent Kit V3 for 2 × 300 bp PE and MiSeq Reagent Kit V2 for 2 × 250 bp PE, respectively.

***Primer sequences***
For the first PCR, taxa-specific universal primers were combined with the MiSeq sequencing primers and six random bases (`N`s) to improve the quality of MiSeq sequencing. For the second PCR, MiSeq adaptor and sequencing primers were combined with index sequences (eight bases denoted by `X` in the following table) to identify each sample. See sample metadata for index sequences of each sample.

Prokaryote 16S, 515F:

TCGTCGGCAGCGTCAGATGTGTATAAGAGACAGNNNNNNGTGYCAGCMGCCGCGGTAA
Prokaryote 16S, 806R:
GTCTCGTGGGCTCGGAGATGTGTATAAGAGACAGNNNNNNGGACTACNVGGGTWTCTAAT

Eukaryote 18S, Euk_1391f:
TCGTCGGCAGCGTCAGATGTGTATAAGAGACAGNNNNNNGTACACACCGCCCGTC
Eukaryote 18S, EukBr:
GTCTCGTGGGCTCGGAGATGTGTATAAGAGACAGNNNNNNTGATCCTTCTGCAGGTTCACCTAC

Fungal ITS, ITS1_F_KYO1:
TCGTCGGCAGCGTCAGATGTGTATAAGAGACAGNNNNNNCTHGGTCATTTAGAGGAASTAA
Fungal ITS, ITS_KYO2:
GTCTCGTGGGCTCGGAGATGTGTATAAGAGACAGNNNNNNTTYRCTRCGTTCTTCATC

Animal COI, mlCOIintF:
ACACTCTTTCCCTACACGACGCTCTTCCGATCTNNNNNNGGWACWGGWTGAACWGTWTAYCCYCC
Animal COI, HCO2198:
GTGACTGGAGTTCAGACGTGTGCTCTTCCGATCTNNNNNNTAAACTTCAGGGTGACCAAAAAATCA

2nd PCR primer for 16S, 18S and ITS, Forward:
AATGATACGGCGACCACCGAGATCTACACXXXXXXXXTCGTCGGCAGCGTCAGATGTGTATAAGAGACAG
2nd PCR primer for 16S, 18S and ITS, Reverse:
CAAGCAGAAGACGGCATACGAGATXXXXXXXXGTCTCGTGGGCTCGGAGATGTGTATAAGAGACAG

2nd PCR primer for COI, Forward:
AATGATACGGCGACCACCGAGATCTACACXXXXXXXXACACTCTTTCCCTACACGACGCTCTTCCGATCT
2nd PCR primer for COI, Reverse:
CAAGCAGAAGACGGCATACGAGATXXXXXXXXGTGACTGGAGTTCAGACGTGTGCTCTTCCGATCT

***Internal standard DNA***

Five artificially designed and synthesized internal standard DNAs, which are similar but not identical to the corresponding region of any existing target organism (e.g., the V4 region of prokaryotic 16S rRNA), were included in the library preparation process to estimate the number of DNA copies (i.e., the quantitative eDNA metabarcoding; Ushio, 2019; Ushio et al., 2018). They were designed to have the same primer-binding regions as those of known existing sequences and conserved regions in the insert region. The sequences were provided in Table S4–S7 in Ushio (2022). Variable regions in the insert region were replaced with random bases so that no known existing sequences had the same sequences as the standard sequences. The numbers of standard DNA copies were adjusted appropriately to obtain a linear regression line between the copy numbers of the standard DNAs and their sequence reads from each sample.
