## Supplementary Figures for "Contrasting ecological communities in rice paddy fields under conventional and no-fertilizer farming practices"

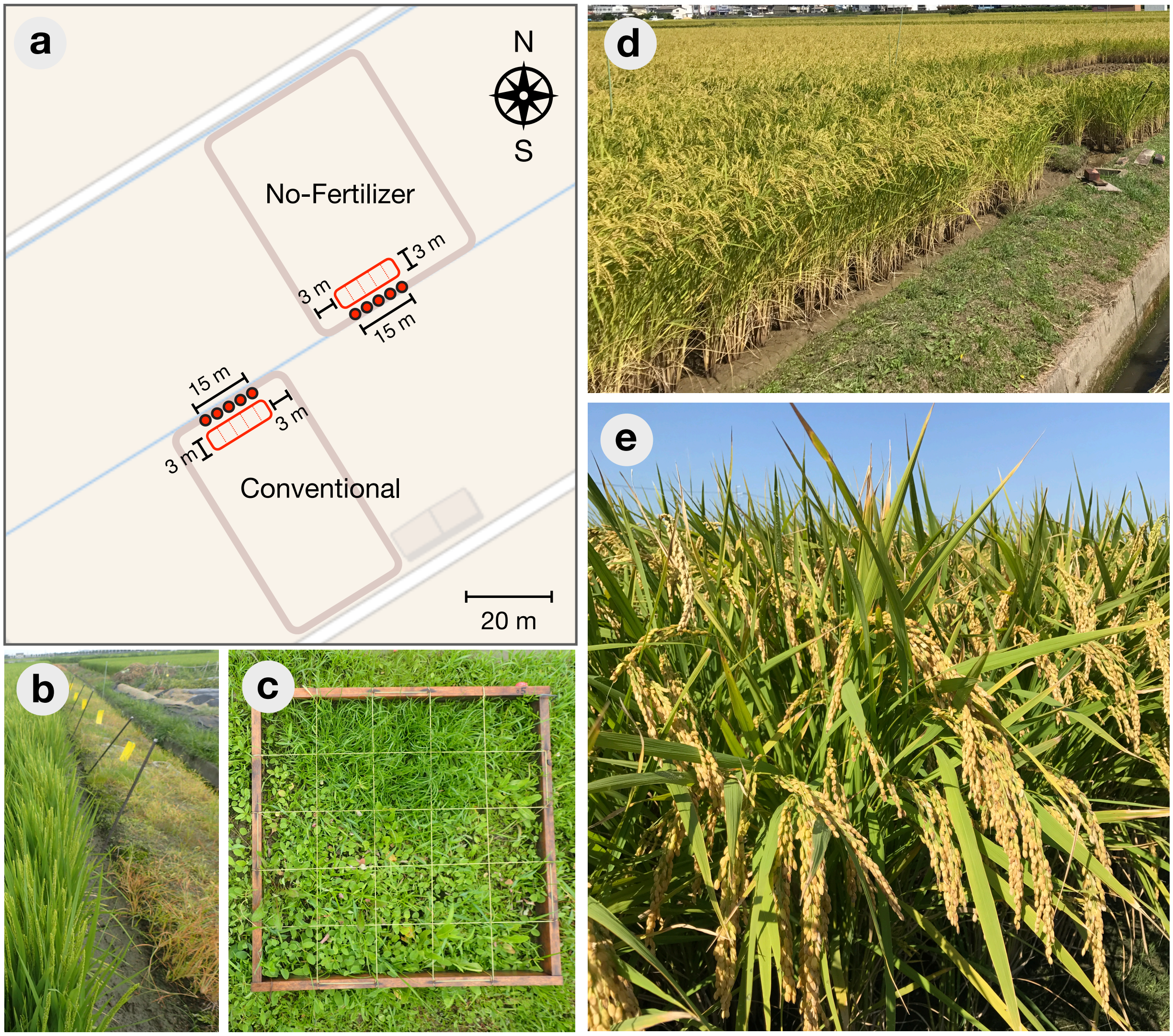


**Figure S1 | Supplementary images from the paddy field survey in 2017**
(**a**) Study site layout including two rice paddy fields: "conventional" and "no-Fertilizer." The two paddy fields are separated by ridges, and we set a 3 m × 15 m plot in each paddy field (indicated by a red rectangle) with a 2-m buffer zone between the plot and the ridge. Each 3 m × 15 m plot was divided into five sections (separated by dashed red lines in each plot), with each section covering an area of 3 m × 3 m. Water samples were collected from all the five sections in each plot to monitor the ecological communities, and the performance of rice individuals were monitored (e.g., rice heights, SPAD, and the numbers of stems and heads) at the same locations. We also set five sampling points on the ridge (indicated by the red circles). We set sticky panels for collecting insects and 50 cm × 50 cm quadrats for monitoring wild weeds. (**b**) Sticky panels for insect monitoring (30 August 2017; ©M. Ushio). (**c**) Plant coverage survey (5 July 2017; ©M. Ushio). Each grid corresponds to 10 cm × 10 cm. (**d**) Yield survey in September 2017 (©S. Yoshinami). (**e**) Rice heads before harvesting (©S. Yoshinami).

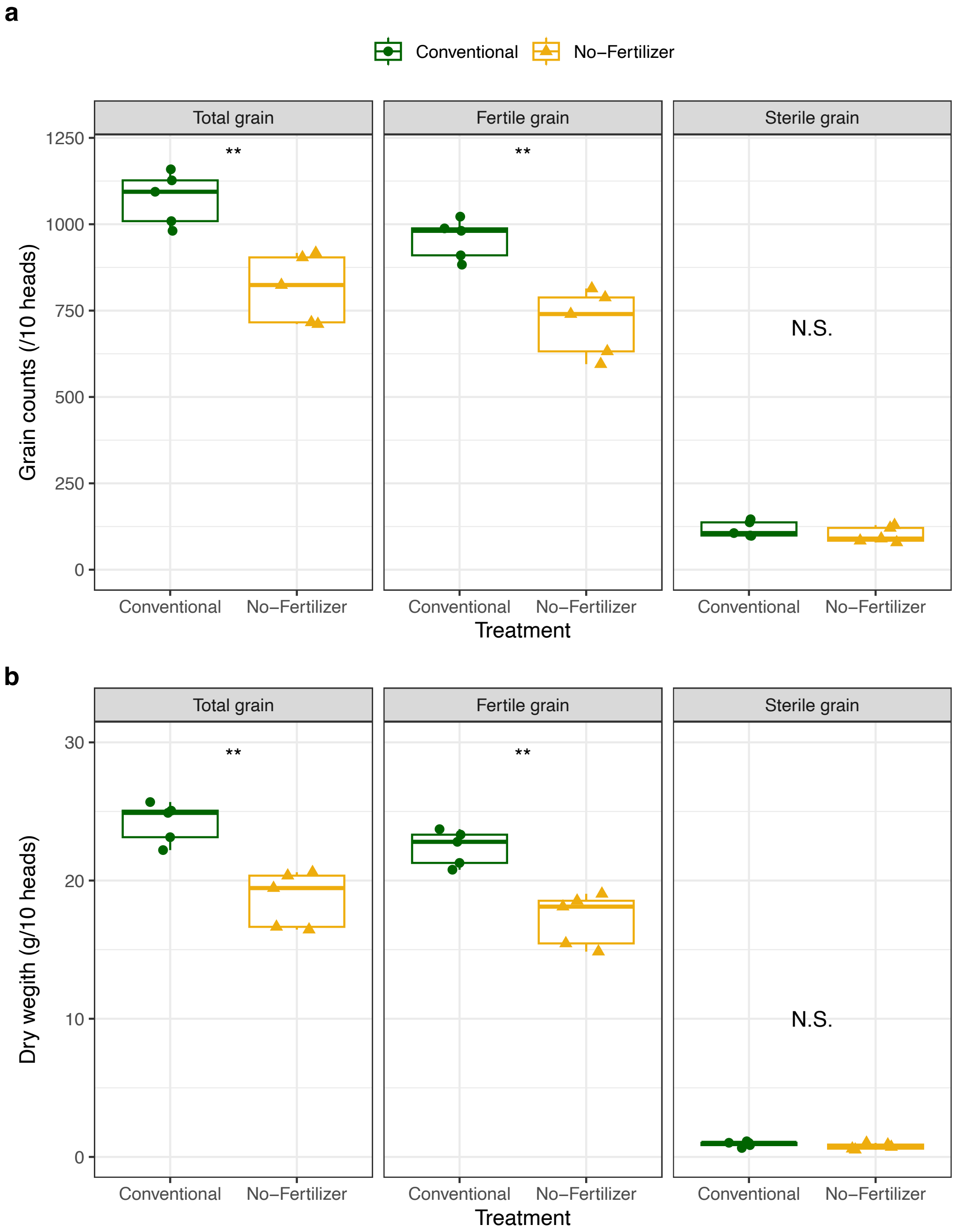


**Figure S2 | Rice grain data**
(**a**) The number of rice grains per 10 rice heads. (**b**) Dry weight (g) of the rice grains per 10 heads. Green and and filled circle indicate data from the conventional paddy fields, and yellow and filled triangle indicate data from the no-fertilizer paddy fields. *$P < 0.05$, **$P < 0.01$, ***$P < 0.001$, and ****$P < 0.0001$. "N.S." indicates that the difference between the two farming practices is not statistically clear.

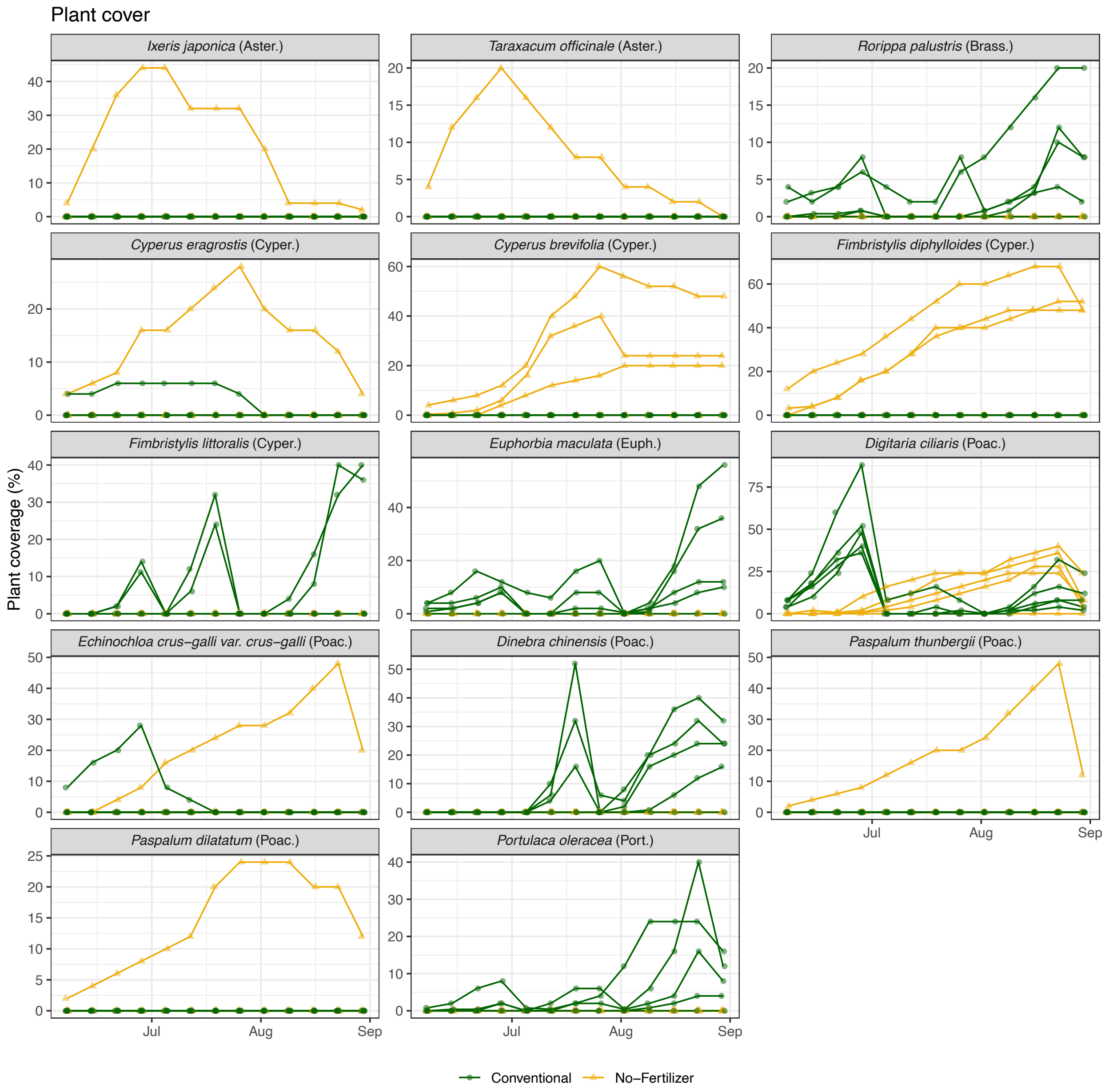


**Figure S3 | The coverage (%) of plant species on the paddy ridges**

Each panel shows the coverage (%) of each plant species as indicated by the strip label. *Taraxacum officinale*, *Cyperus eragrostis*, *Euphorbia maculata*, and *Paspalum dilatatum* are alien species, while the other species are native species. *Rorippa palustris*, *Fimbristylis littoralis*, *Euphorbia maculata*, *Digitaria ciliaris*, *Echinochloa crus-galli* var. *crus-galli*, *Dinebra chinensis*, and *Portulaca oleracea* are annual species, while the other plant species are perennial species. Solid lines indicate mean values. Green and and filled circle indicate data from the conventional paddy field, and yellow and filled triangle indicate data from the no-fertilizer paddy field.

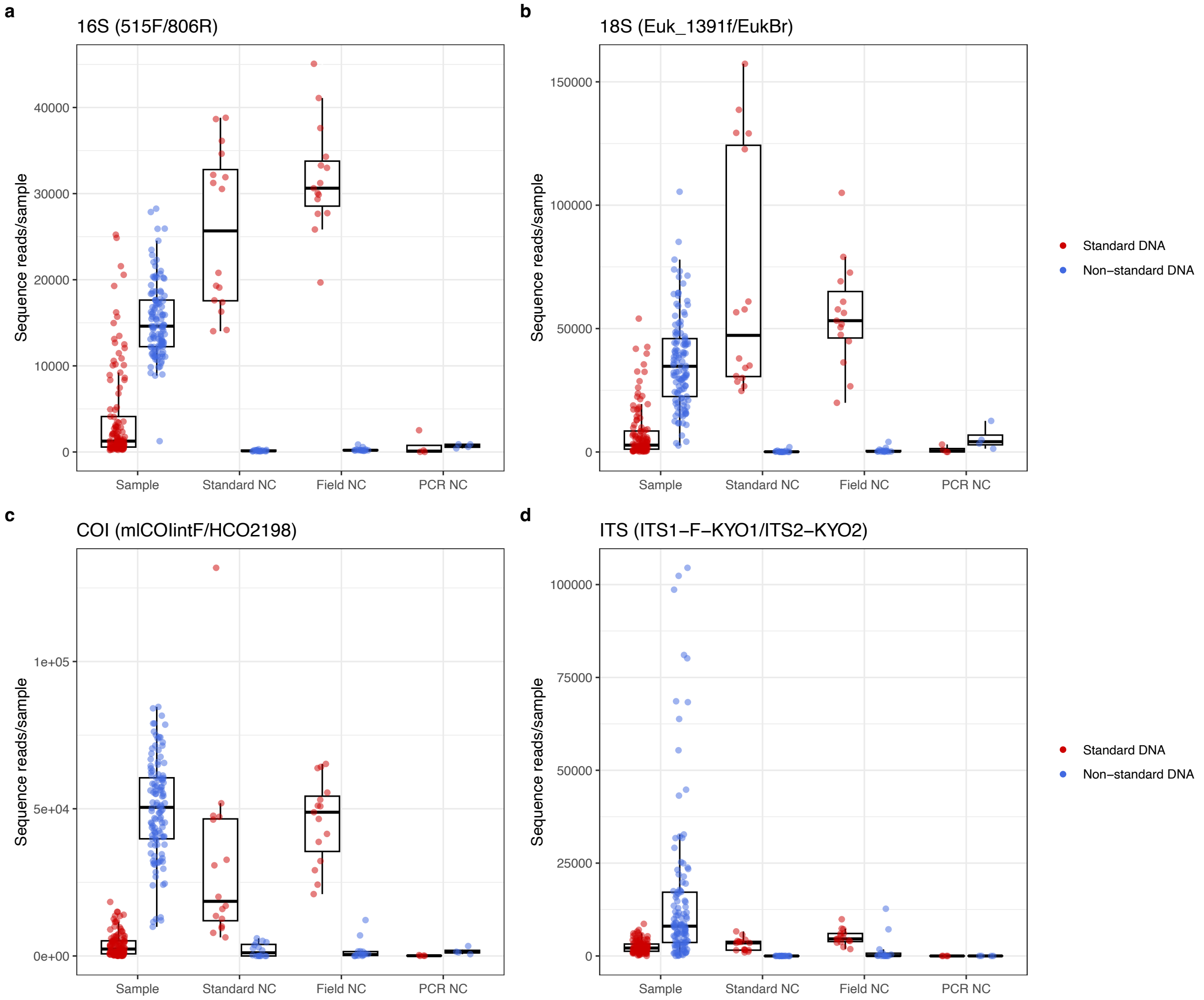


**Figure S4 | Summary of Sequence reads from the quantitative eDNA metabarcoding**
Sequence reads of standard DNA and non-standard DNA (real organism DNA) for (**a**) 16S, (**b**) 18S, (**c**) COI, and (**d**) ITS regions. "Sample," "Standard NC," "Field NC," and "PCR NC" indicate real samples, standard DNA negative controls (NC) (MilliQ water with standard DNA only), field NC (MilliQ water collected on site and processed with standard DNA) and PCR NC (MilliQ water included in the PCR without standard DNA), respectively. Red and blue points indicate sequence reads of standard DNA and non-standard DNA detected from each sample, respectively.

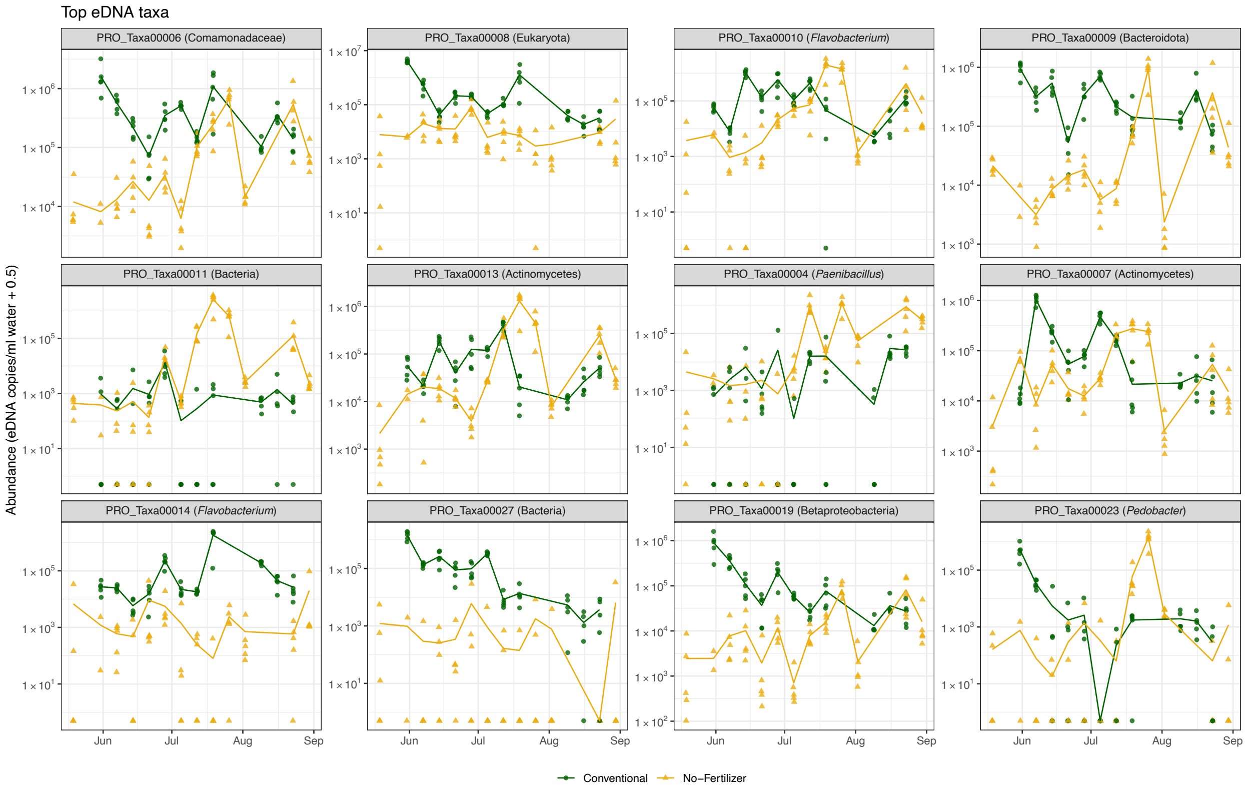


**Figure S5 | Top 12 most abundant taxa detected by the quantitative eDNA metabarcoding**
The top 12 eDNA taxa based on the total eDNA copy numbers. Green and and filled circle indicate data from the conventional paddy fields, and yellow and filled triangle indicate data from the no-fertilizer paddy fields. The solid line indicate the mean eDNA copy numbers for each paddy field. Strip labels indicate the taxa code and taxa name assigned by Claident.

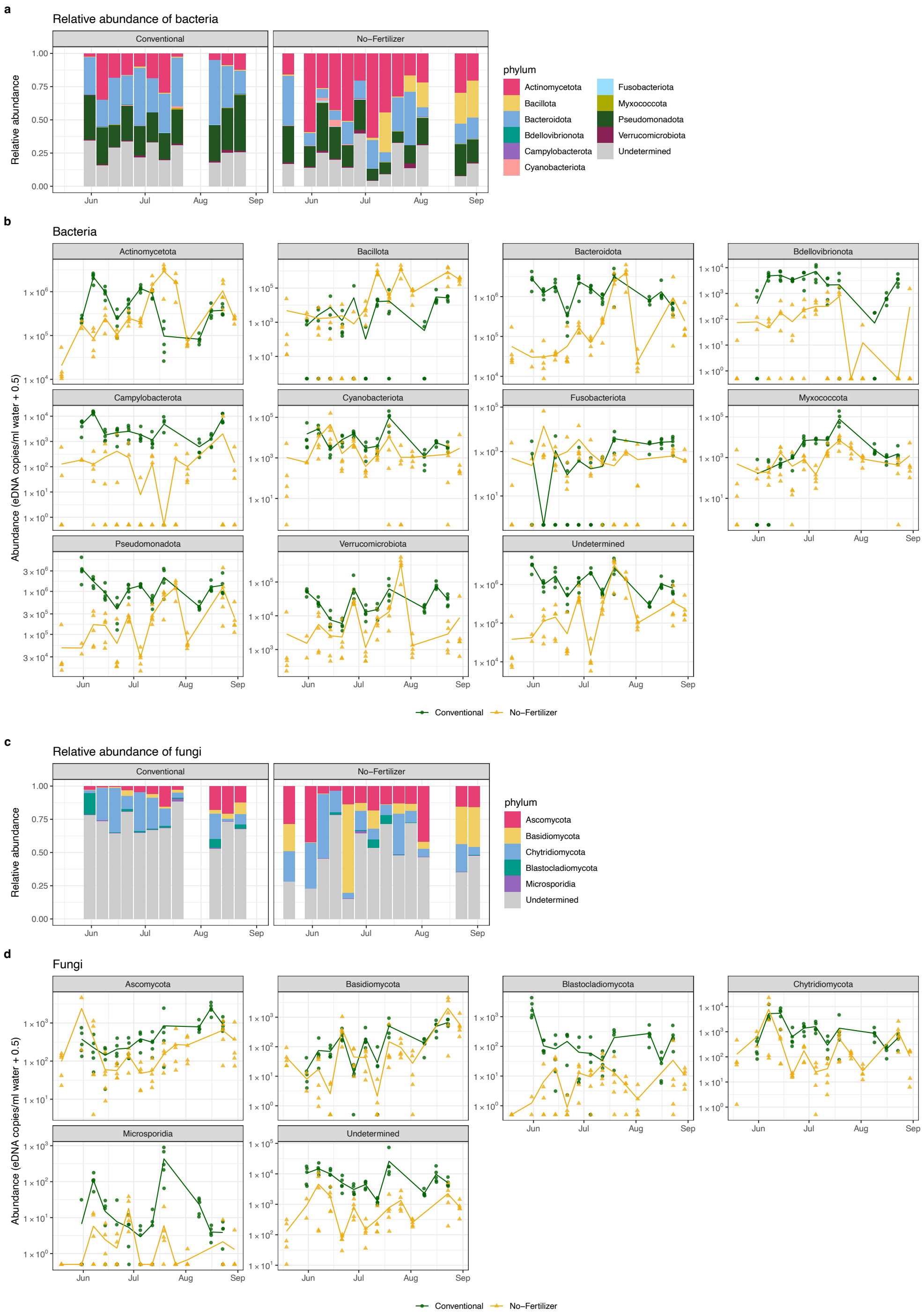


**Figure S6 | Temporal dynamics of major bacterial and fungal phyla detected by the quantitative eDNA metabarcoding**

(**a**) The relative and (**b**) absolute abundance of major phyla of bacteria, and (**c**, **d**) those of fungi. Green and and filled circle indicate data from the conventional paddy fields, and yellow and filled triangle indicate data from the no-fertilizer paddy fields. The solid line indicate the mean eDNA copy numbers for each paddy field. Strip labels indicate the taxa code and taxa name assigned by Claident.

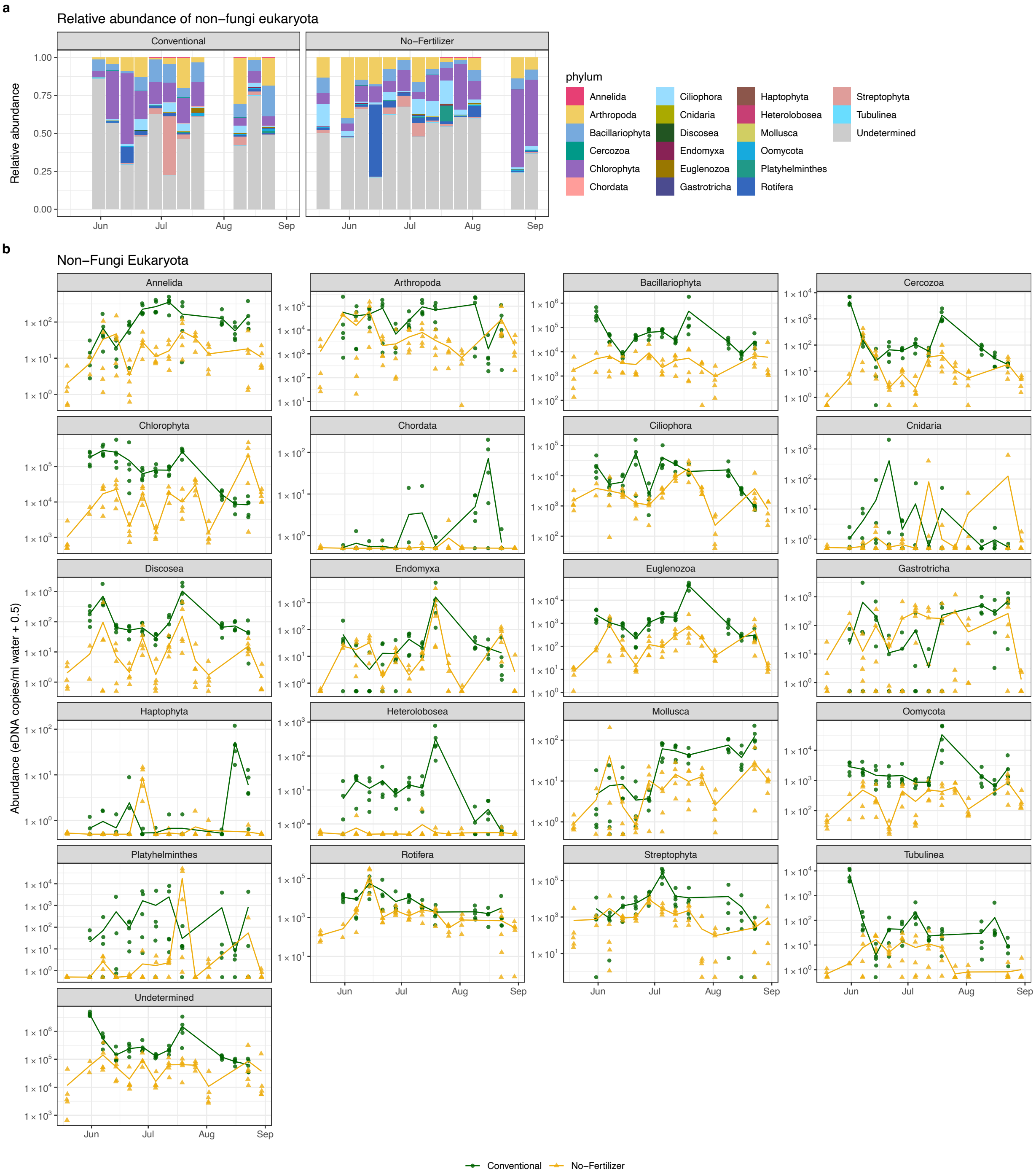


**Figure S7 | Temporal dynamics of major non-fungal eukaryota detected by the quantitative eDNA metabarcoding**

(**a**) The relative and (**b**) absolute abundance of major phyla of non-fungi eukaryota. Green and and filled circle indicate data from the conventional-farming paddy fields, and yellow and filled triangle indicate data from the no-fertilizer paddy fields. The solid line indicate the mean eDNA copy numbers for each paddy field. Strip labels indicate the taxa code and taxa name assigned by Claident.

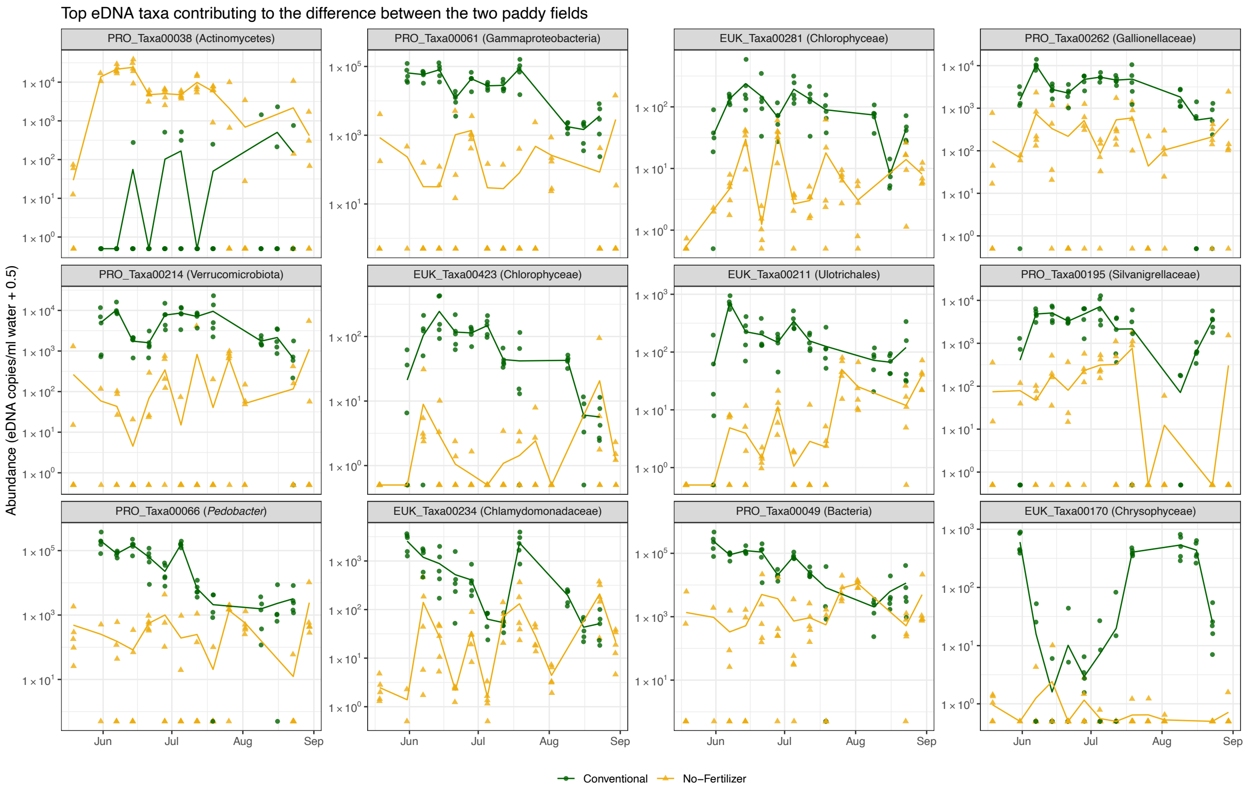


**Figure S8 | eDNA taxa that differentiate the conventional and no-fertilizer paddy fields**
The top 12 eDNA taxa based on their contribution to the differences in community compositions between the two paddy fields. Green and and filled circle indicate data from the conventional paddy fields, and yellow and filled triangle indicate data from the no-fertilizer paddy fields. The solid line indicate the mean eDNA copy numbers for each paddy field. Strip labels indicate the taxa code and taxa name assigned by Claident.

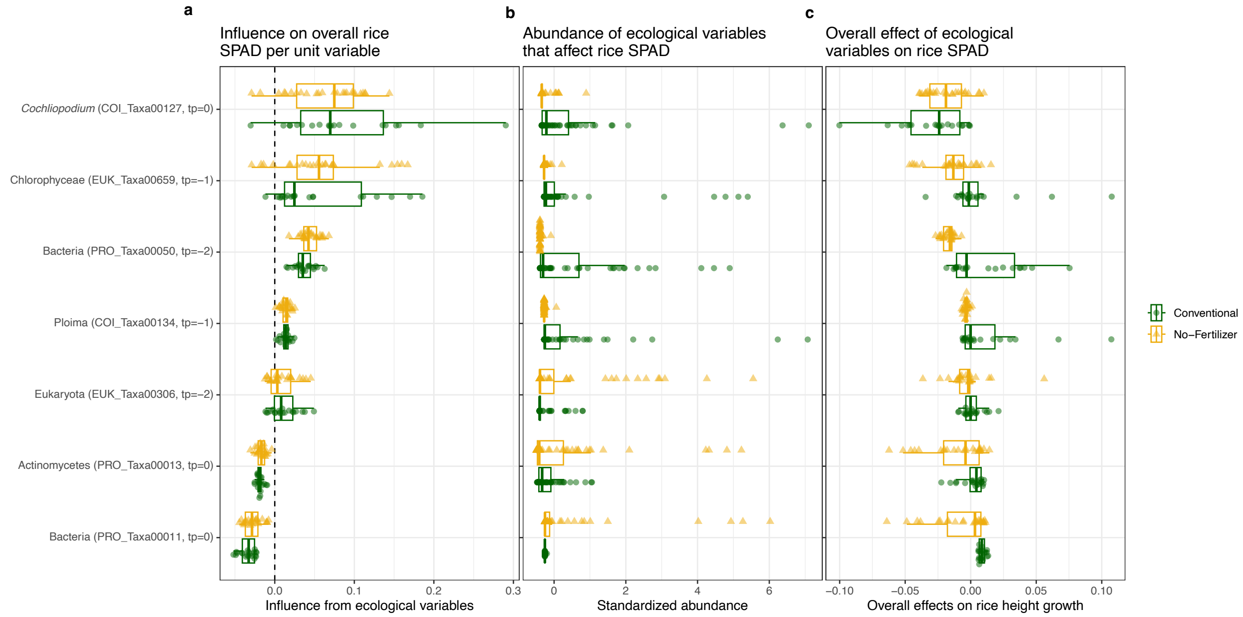


**Figure S9 | Influences of ecological variables on the SPAD values in the conventional and no-fertilizer paddy fields**

Each panel shows (**a**) the influence of ecological variables on the rice SPAD values per unit variable (i.e., coefficients of the MDR S-map), (**b**) the abundance of ecological variables, and (**c**) the overall effects of ecological variables on rice height growth (calculated by "the MDR S-map coefficient $\times$ Abundance"), respectively. Green and and filled circle indicate data from the conventional paddy fields, and yellow and filled triangle indicate data from the no-fertilizers paddy fields.

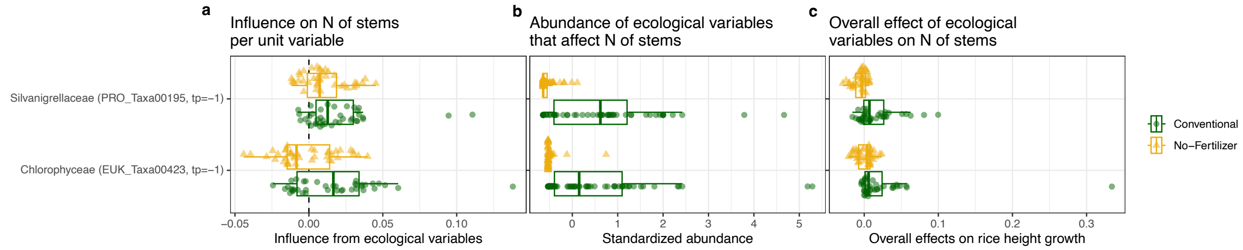


**Figure S10 | Influences of ecological variables on the number of rice stems in the conventional and no-fertilizer paddy fields**

Each panel indicates (**a**) the influence of ecological variables on the number of rice stems per unit variable (i.e., coefficients of the MDR S-map), (**b**) the abundance of ecological variables, and (**c**) the overall effects of ecological variables on rice height growth (calculated by "the MDR S-map coefficient $\times$ Abundance"), respectively. Green and and filled circle indicate data from the conventional paddy fields, and yellow and filled triangle indicate data from the no-fertilizers paddy fields.

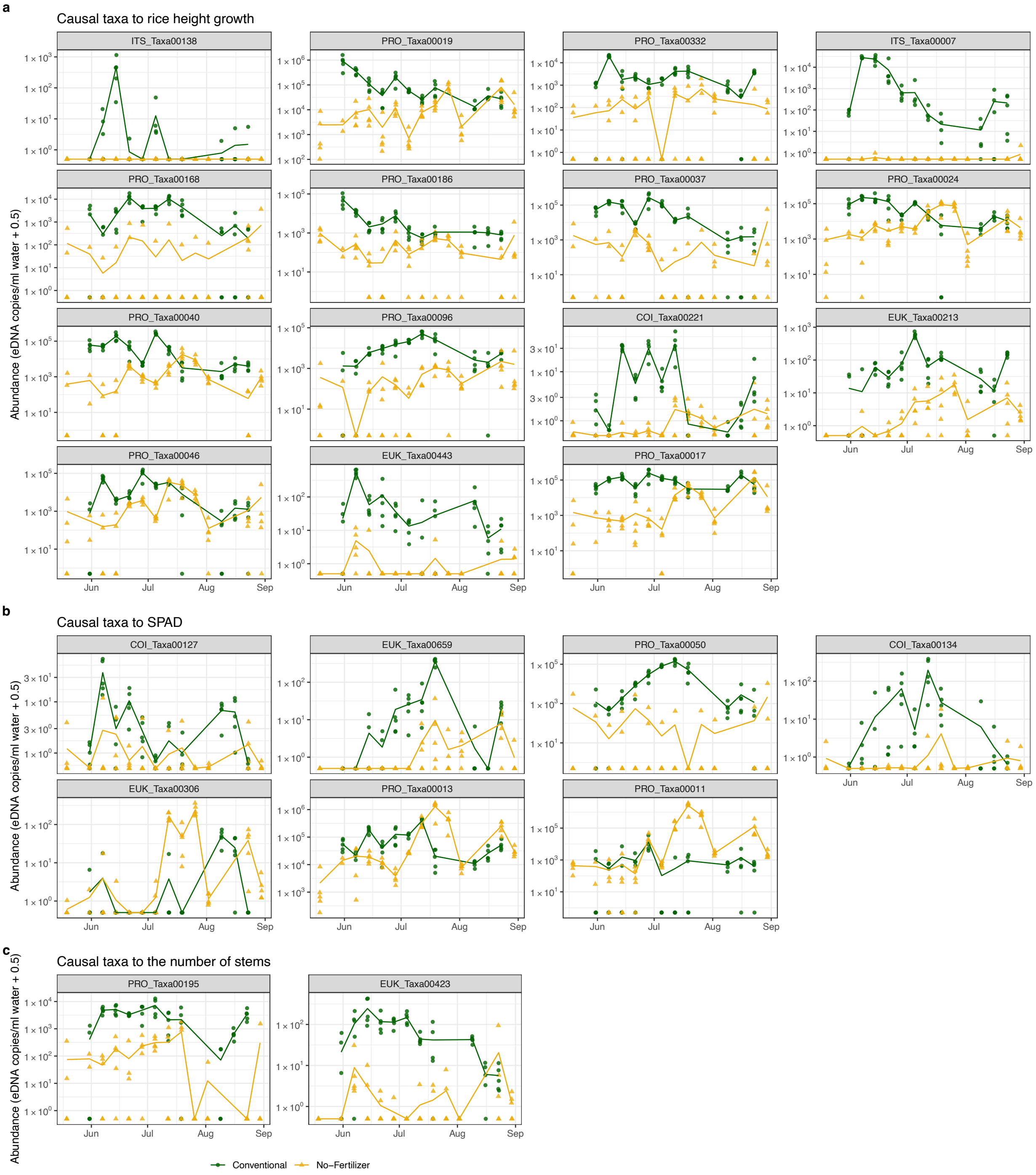


**Figure S11 | Temporal dynamics of taxa that causally influenced rice performance**
Temporal dynamics of (**a**) taxa that causally influenced the rice height growth, (**b**) SPAD values, and (**c**) the number of rice stems. Green and and filled circle indicate data from the conventional paddy fields, and yellow and filled triangle indicate data from the no-fertilizers paddy fields.
